# Beyond D_max_ and DC_50_: A simple tool to evaluate the complex PROTAC-mediated kinetic degradation dynamics

**DOI:** 10.64898/2026.09.18.752745

**Authors:** Smitha Kizhake, Amarnath Natarajan

## Abstract

Protein degradation mediated by PROteolysis TArgeting Chimeras (PROTACs) is a complex, dynamic, and multi-step process. The structural diversity of PROTACs, coupled with cellular heterogeneity, introduces considerable variability at each stage. Therefore, a straightforward yet comprehensive approach is essential to effectively dissect these mechanisms. The existing methods for evaluating PROTAC degradation are often time-consuming, labor-intensive, or expensive, and many capture only a partial view of the entire degradation process. Consequently, multiple complementary assays are typically required to obtain the complete picture. Here, we use commercially available Cyclin dependent Kinase 6 (CDK6) targeting PROTACs to demonstrate that a simple, single-step, no-wash, additive-free, GFP-based live-cell imaging approach can rapidly generate dynamic degradation metrics (D_max_, R_max_, TD_max_, CD_max_, DC_50_, K_m_, T_max_, K_deg_, V_max_, DT_50_, *τ* and TR_max_) for comparing and ranking PROTACs within a controlled reporter system. The rapid assay turnaround time, excellent assay qualities and compatibility with high-throughput (HT) formats will facilitate quick hit identification and serve as a guiding assay for hit-to-lead optimization campaigns.

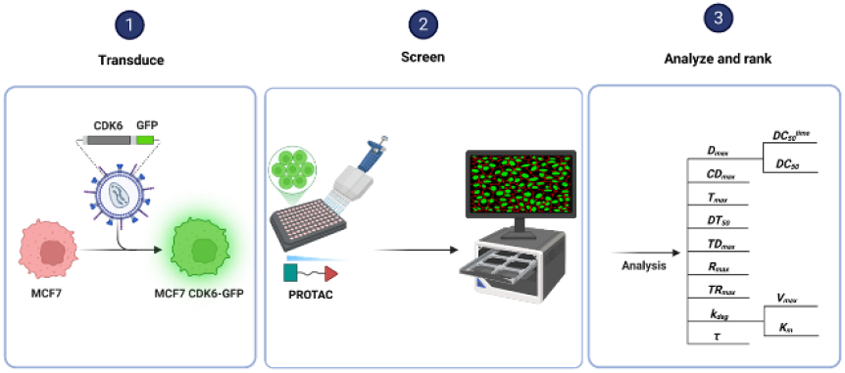

## Introduction

PROTACs are heterobifunctional molecules that hijack the endogenous cellular degradation machinery^1^ and have been the hotbed of research for over two decades.^2-9^ From the first peptide-based PROTAC in 2001 to the first FDA-approved PROTAC in 2026, the field has advanced remarkably.^10-14^ However, the complexity of cellular systems, combined with the intricacy of PROTAC degradation mechanism, has posed significant challenges along its developmental path.^15-22^ PROTAC-mediated protein degradation is a multi-step process involving - cell permeation, target engagement, through binary complex (BC) and ternary complex (TC) formation, primary ubiquitination, ubiquitin chain elongation and ultimately proteasomal degradation. Each of these steps is in turn, cell-type dependent, and any one of them could be the rate-limiting step. *In vitro* and *in vivo* assays^23^ (Fig. 1) and mathematical models^24-27^ have been developed to gain a better understanding of the degradation process.

**Fig. 1.**
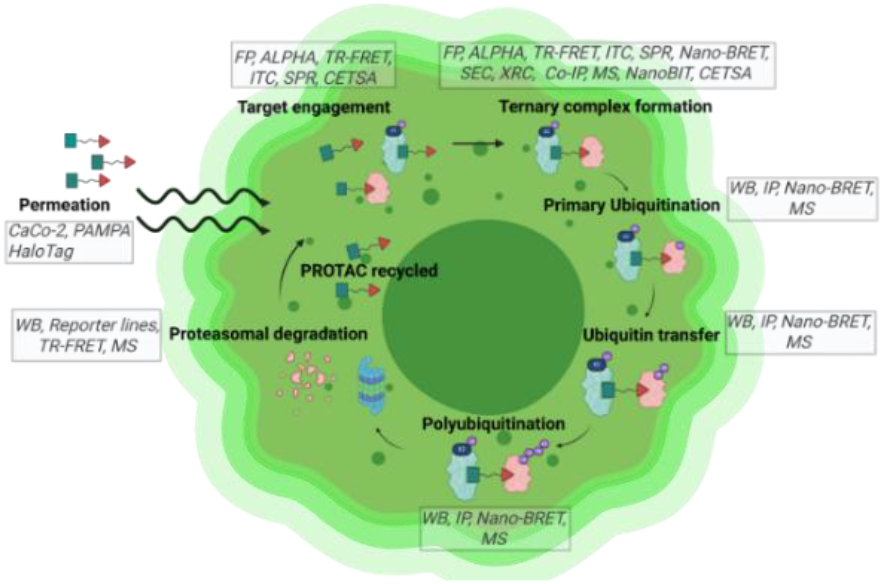
Pictorial representation of the PROTAC degradation process and the associated assays used for development.

Many of the *in vitro* biophysical assays like Fluorescence Polarization (FP),^28-33^ Amplified Luminescent Proximity Homogeneous Assay (ALPHA),^34-37^ Time-Resolved Fluorescence Energy Transfer (TR-FRET),^38-41^ Isothermal Titration Calorimetry (ITC),^29, 34, 42-44^ Surface Plasma Resonance (SPR),^32, 35, 36, 44, 45^ Size Exclusion Chromatography (SEC) ^42, 46, 47^ and Crystallography,^29, 31, 34, 48-50^ are either time-consuming, labor-intensive, expensive or not HT amenable. Moreover, they provide a limited, static, snapshot of the PROTAC-mediated degradation process and fail to capture the kinetic information that is critical in a dynamic cellular environment. Furthermore, because these assays are typically performed under non-physiological conditions, caution should be exercised when extrapolating results from biophysical assays to cellular systems, as recombinant systems cannot fully recapitulate the complexity of the cellular environment^51^

While cell-based *in vitro* assays like, Nano-Bioluminescence Resonance Energy Transfer (Nano-BRET),^37, 52-58^ Co-ImmunoPrecipitation (Co-IP),^31, 36, 59-61^ NanoLuc Binary Technology (NanoBiT),^56, 57, 60, 61^ Mass Spectrometry (MS),^31, 36, 37, 62-68^ Cellular Thermal Shift (CETSA),^36, 55, 58, 69^ Western Blot (WB),^30, 35, 70-73^ GFP-fusion protein reporter^10, 74-77^ and High-Affinity Binary Technology (HiBiT)^54, 78, 79^ assays are more versatile and offer valuable insights into the degradation process, they are typically not comprehensive. Recently, HiBiT based live-cell assays have been widely used to obtain PROTAC mediated kinetic degradation data. While the HiBiT is an attractive platform with many advantages, the reliance on an exogenous substrate to measure assay readouts leads to increased long-term assay costs. Therefore, a rapid, simple, and cost-effective assay is needed, that enables comprehensive characterization of PROTAC degradation kinetics while accelerating the discovery and development process.

CDK6, an important cell cycle regulatory protein,^80^ is frequently upregulated in cancers and the upregulation leads to disease progression.^81^ Overexpression of CDK6 in breast cancer contributes to its resistance against inhibitors,^82^ which can be overcome by employing PROTACs.^83,84^ Green Fluorescence Protein (GFP) was first discovered serendipitously in 1962^85^ and has subsequently revolutionized our ability to study many cellular processes. The ability of GFP to withstand photobleaching^86^ coupled with the fact that GFP-tagged proteins display similar degradation profiles to endogenously expressed proteins,^87^ provides a strong rationale to employ GFP-fused PoI’s to study PROTAC degradation.

We evaluated a panel of cancer cell lines, for CDK6 expression, and identified MCF7 as having undetectable endogenous CDK6 protein levels, making it an ideal choice to introduce GFP-tagged CDK6 and generate an MCF7 CDK6-GFP cell line for use as a controlled reporter platform. The cell line was validated by assessing the phosphorylation of Rb(S^780^) by immunoblotting and by the release of the G1 checkpoint by cell cycle analysis. Subsequently we demonstrated how the degradation kinetics of 10 commercially available CDK6 PROTACs [BSJ-02-162,^88^ BSJ-03-123,^88^ BSJ-03-204,^88^ BSJ-04-132,^88^ CDK4/6 Degrader I,^89^ CP-10,^90^ RSS0680,^91^ XY028-133,^92^ XY028-140 (MS-140)^93^ and YX-2-107^94^] was obtained in a single-step, no-wash, additive-free, live-cell GFP imaging assay performed under physiological conditions throughout the experiment. The rapid generation of the stable cell line combined with the short assay duration, enabled fast experimental turnaround. Furthermore, this approach was cost-effective, as it did not require an exogenous substrate, making it a practical live-cell assay for monitoring PROTAC degradation. Importantly, an orthogonal assay of endogenous CDK6 degradation in BxPC3 cells showed high correlation with the CDK6-GFP degradation seen in the imaging studies, supporting the fact that the GFP loss reflects CDK6 depletion.

PROTACs can be ranked based on three criteria (Fig. 2):

**Fig. 2.**
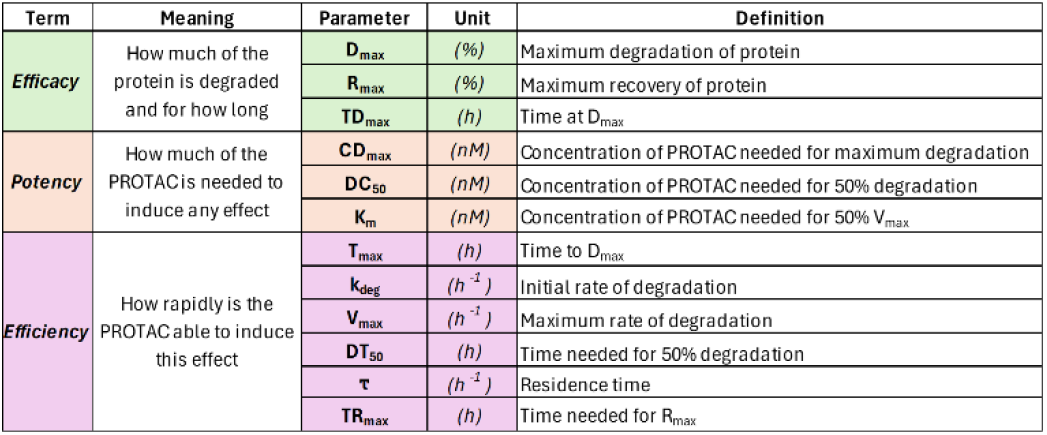
Degradation parameters obtained from the screen with their acronyms and definitions.

- **Efficacy** (*extent and duration of PoI degradation*) characterized by the maximum degradation of the PoI (**D**_**max**_),^95^ maximum recovery of the PoI (**R**_**max**_)^96^ and duration at D_max_ (**TD**_**max**_)^54^
- **Potency** (*concentration of PROTAC required to induce degradation*) characterized by concentration at which maximum degradation occurs (**CD**_**max**_), concentration required for 50% degradation of the PoI (**DC**_**50**_)^95^ and the concentration needed to reach 50% of the maximum degradation rate (**K**_**m**_)^97^
- **Efficiency** (*rate of degradation of PoI by the PROTAC*) characterized by the time to reach maximum degradation (**T**_**max**_),^96^ time required for 50% degradation of the PoI (**DT**_**50**_),^98^ initial degradation rate (**k**_**deg**_),^96^ maximum degradation rate (**V**_**max**_),^97^ residence time (***τ***) and time to maximum recovery (**TR**_**max**_).

Here, using the MCF7 CDK6-GFP cell line and commercially available CDK6 PROTACs, we demonstrate how all the above parameters can be obtained from two screens. The PROTACs were subsequently ranked based on their individual performance metrics (**efficacy, potency**, and **efficiency**) and their overall performance across all three criteria, which can serve as a valuable tool in hit-to-lead optimization.

## Results

### Cell line validation

MCF7 Wild type (WT) cells were transduced at varying multiplicities of infection (MOI) (Fig. 3) and the resulting clones showed similar morphology (Fig. S1A) and growth patterns (Fig. S1B) to the WT cells. Subsequently, Flow Cytometry (Fig. S1C) and WB were done to check for variations in cell cycle and CDK6-GFP levels, respectively. The clones showing equivalent amounts of CDK6 and GFP protein expression levels (5F, 10R, 2.5R, 1.25R) (Fig. S1D) were selected and subjected to an initial degradation assay with a CDK6 PROTAC (BSJ-03-123) to identify the optimal clone for all subsequent assays (Fig. S1E). Based on the overall degradation (Fig. S1F), clone 1.25R was then selected for all further experiments. The relative mRNA levels of the key cell cycle proteins, E3 Ligases (E3L), and housekeeping proteins in MCF7 WT are shown in Fig. S1G (www.proteinatlas.org). Comparison of the baseline protein levels in WT and 1.25R by immunoblotting (Fig. S1H) showed that neither the E3Ls (Cereblon - CRBN, Von-Hippel Lindau - VHL) nor the housekeeping proteins (β-Actin, GAPDH, HSP90, Tubulin and Vinculin) were altered by the transduction. The elevated p-Rb (S^780^) levels observed in the CDK6-GFP line compared to CDK6-WT resulted in the release of the G1 checkpoint as previously reported,^82^ and validated the MCF7 CDK6-GFP cell line.

**Fig. 3.**
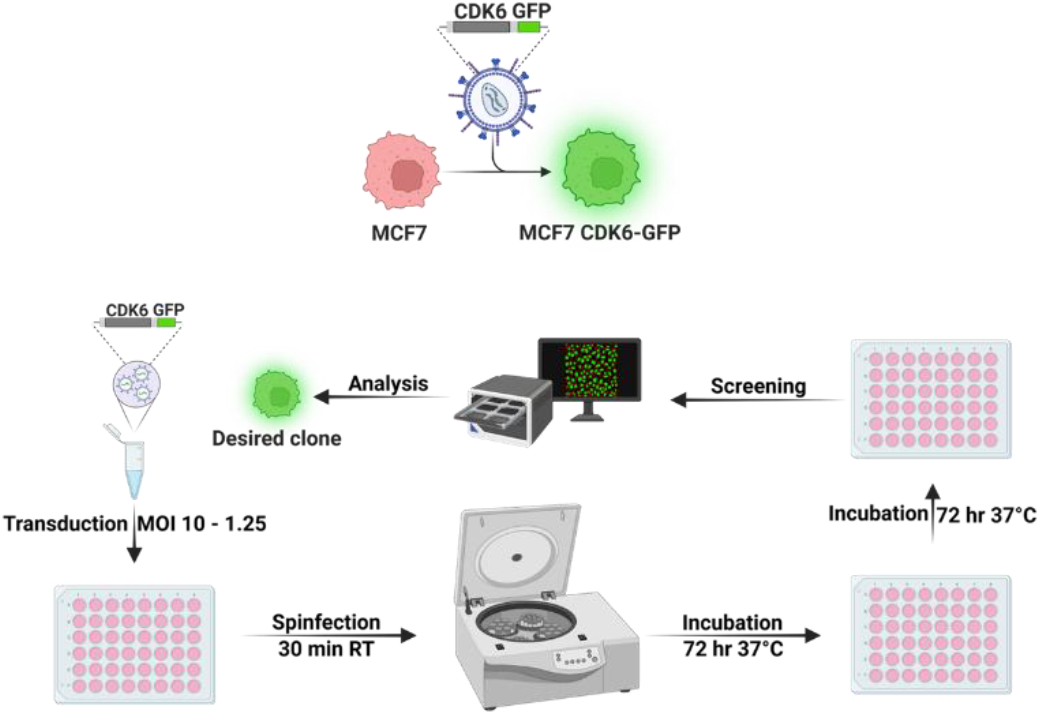
Schematic for the generation and selection of the desired MCF7 CDK6-GFP clone.

### Assay validation

The validated cell line was subjected to assay validation by evaluating the following parameters^99^ –

- **Noise** (fluctuations in the signal)
- **Linearity** (proportionality of signal to amount of sample)
- **Limit of Detection** (LoD – the lowest amount of sample that is distinguishable from the background)
- **Limit of Quantitation** (LoQ - the lowest amount of sample that can be meaningfully measured)
- **Dynamic range** (window of measurement)
- **Precision** (reproducibility of sample measurements)
- **Reproducibility** (precision of sample measurement across varied conditions)
- **Sensitivity** (ability to measure minor changes)
- **Specificity** (ability to distinguish a sample for a non-target)
- **Selectivity** (ability to distinguish a sample from a closely related member).
- **Robustness** (reliability in sample values despite slight changes)

Assay validation was performed in two stages. Primary assay validation (noise, linearity, LoD, LoQ, dynamic range, precision, reproducibility, sensitivity, specificity, and selectivity) was done to optimize assay conditions and Secondary assay validation (robustness) was done to demonstrate assay translatability.

The assay noise showed low variability (%CV – 9%) (Fig. 4A) and the assay was linear (R^2^ – 0.99) over 2 orders of magnitude (Fig. 4B) of cell densities indicating a good dynamic range. The linear fit of this data was used to determine the LoD and LoQ for this assay. The LoD for the assay was 400 cells/well and the LoQ was for 1200 cells/well, for a 96-well plate (Fig. 4C). Applying a ± 10% tolerance relative to the 0% DMSO reading, we found that the maximum tolerated DMSO concentration over a 72-hour period was 1% (Fig. 4D). The low inter-day assay variance (%CV – 6%) (Fig. 4E) attested to the precision while the low inter-plate and intra-day variability (%CV – 8%) confirmed the reproducibility of the assay (Fig. 4F). The high Signal-to-noise (S/N) (> 1000) combined with the low LoD indicated that the assay had good sensitivity.

**Fig. 4.**
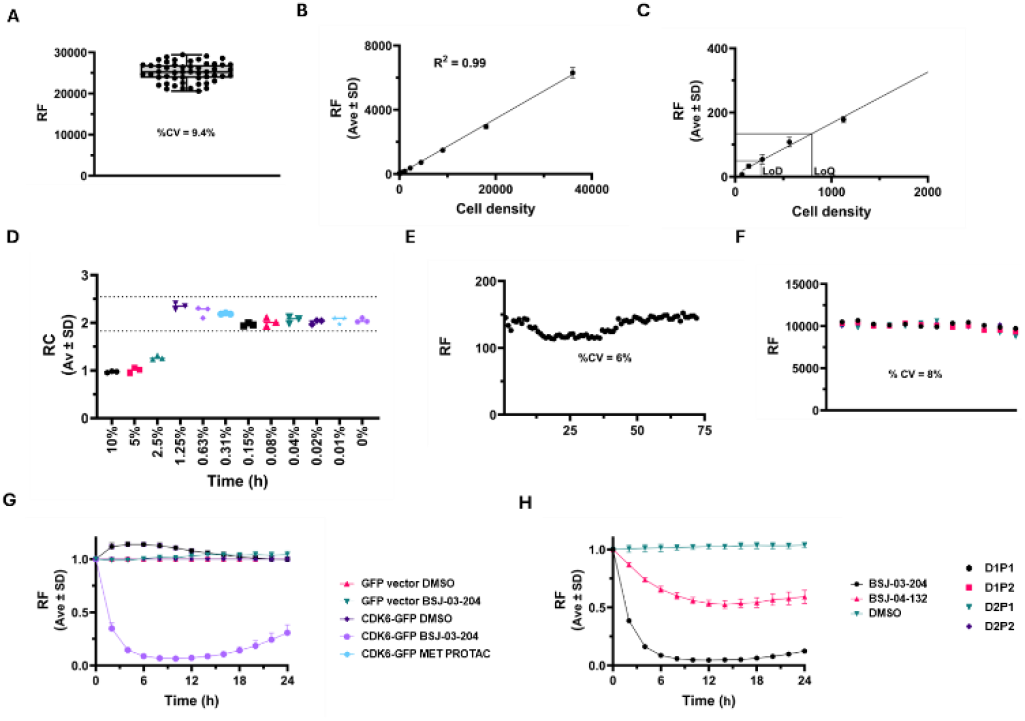
Assay validation - A) Noise determined by imaging 60 wells containing MCF7 CDK6-GFP cells B) Linearity measured by serial dilution of MCF7 CDK6-GFP cells C) Determination of LoD and LoQ D) DMSO tolerance over a 72-hour period E) Precision and reproducibility measured by imaging cells treated with 0.1% DMSO F) Inter and intra-day variability measured by imaging cells treated with 0.1% DMSO G) Specificity determined by treating cells with 1 µM BSJ-03-204 or MET PROTAC H) Selectivity determined by treating cells with 1 µM BSJ-03-204 or BSJ-04-132.

Comparison of CDK6 degradation by BSJ-03-204 (CDK6 PROTAC) and a MET PROTAC (previously published in our lab)^98^ in the MCF7-CDK6-GFP and GFP-vector line cell lines showed selective degradation only for the CDK6 PROTAC in the MCF7 CDK6-GFP line (Fig. 4G). The ability of the MCF7 CDK6-GFP line to distinguish between two closely related PROTACs – BSJ-03-204 (Palbociclib based) and BSJ-04-132 (Ribociclib based). demonstrated assay specificity (Fig. 4H).

### Degradation profile

Following cell line and assay validation, the CDK6 PROTACs were screened (Fig. 5A and Fig. 5B) to derive kinetic parameters. RSS0680, a pan-kinase PROTAC, was included due to its reported off-target degradation of CDK6^91^ and BSJ-04-132, a CDK4-selective PROTAC, was included as another control.

**Fig. 5.**
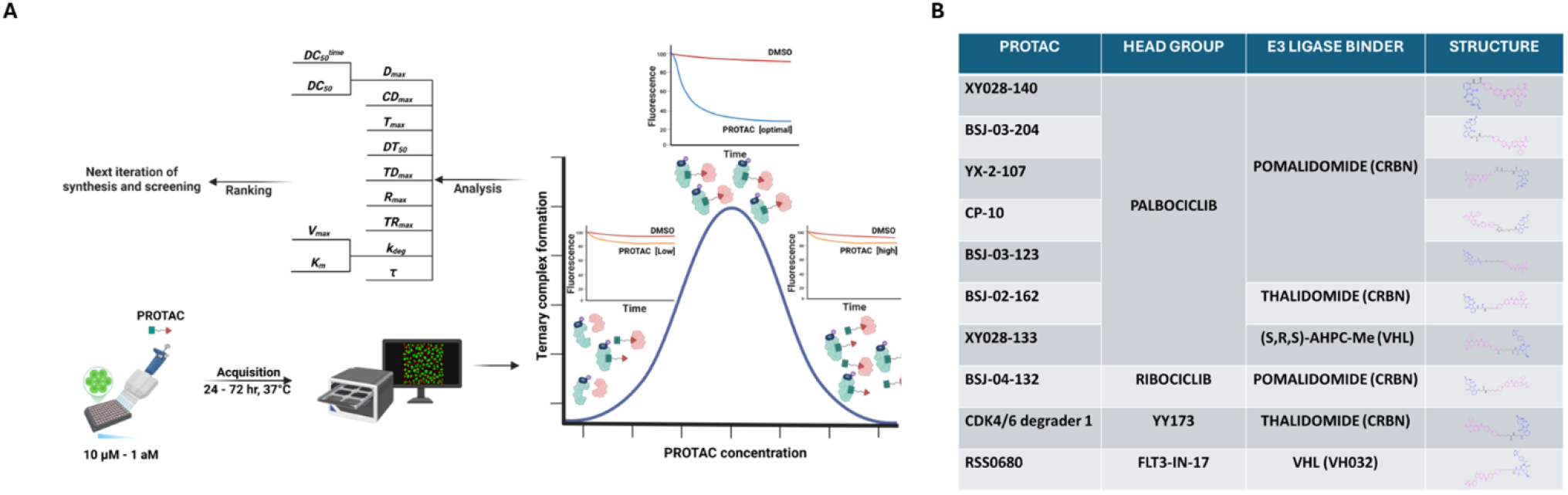
PROTAC screen A) Schematic of the PROTAC screen showing the calculated parameters B) CDK6 PROTACs used in this study.

Results from a 24-hour screen are shown in Figs. 6A - K.

**Fig. 6.**
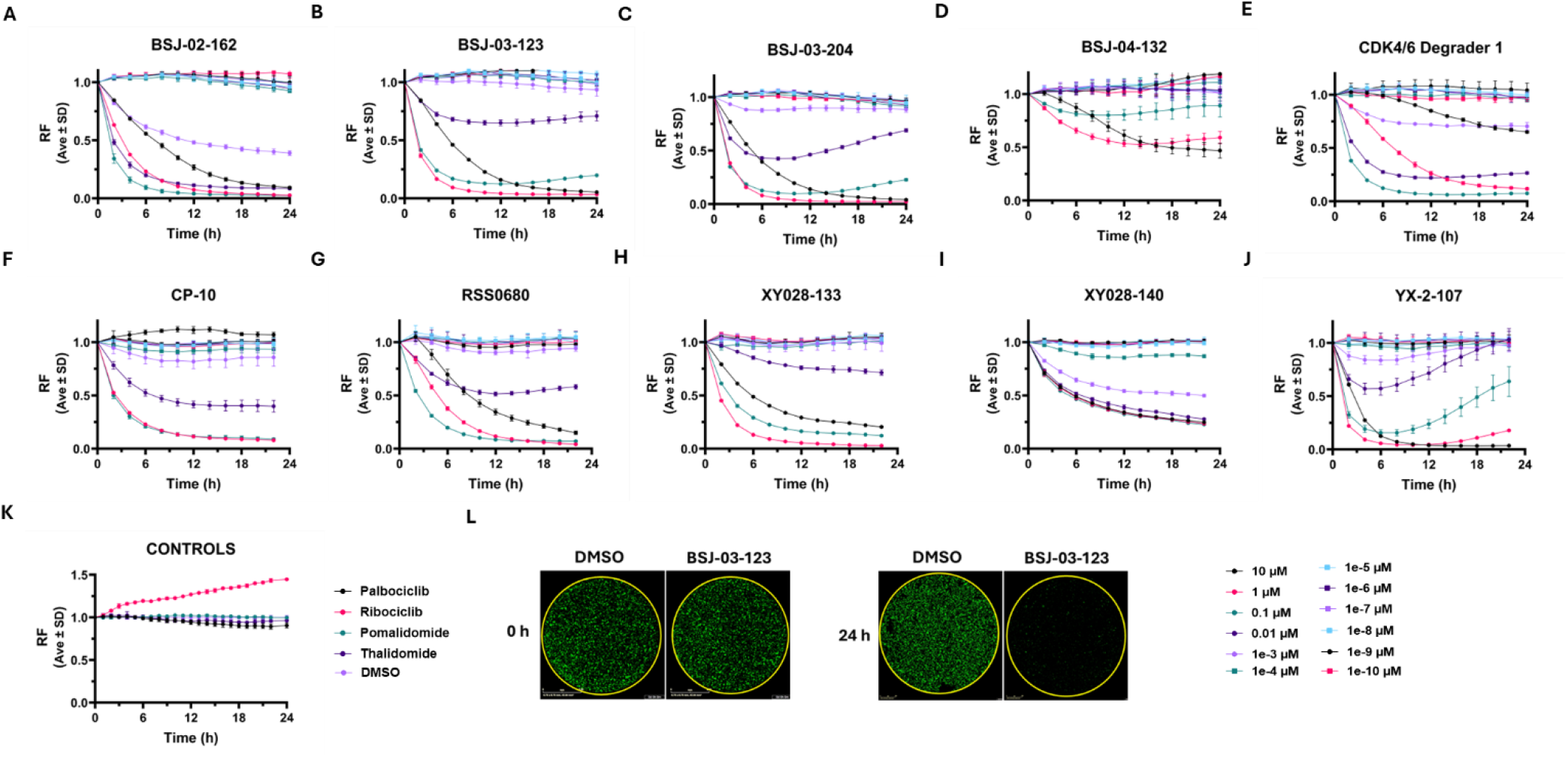
Dose-response curves of all PROTACs A) BSJ-02-162 B) BSJ-03-123 C) BSJ-03-204 D) BSJ-04-132 E) CDK4/6 degrader 1 F) CP-10 G) RSS0680 H) XY028-133 I) XY028-140 J) YX-2-107 K) Controls L) Time lapse image showing the degradation of CDK6-GFP by a PROTAC compared to DMSO (*image contrast digitally enhanced to aid visualization*).

A representative image showing CDK6-GFP degradation in cells treated with either a PROTAC or DMSO, for 24-hours, is shown in Fig. 6L. The 24-hour time lapse degradation videos for all the PROTACs are shown in Fig. S2.CLE

### Parameters obtained from degradation curve

A representative plot with the parameters obtained from the 24-hour degradation curve (**D**_**max**_, **T**_**max**_, **CD**_**max**_, **k**_**deg**_ and **DT**_**50**_) is shown in Fig. 7A. Figs. S3A – I show the curves for all the PROTACs. The amount of CDK6 degraded can be calculated from D_max_ value as shown below^78^

**Figure 7.**
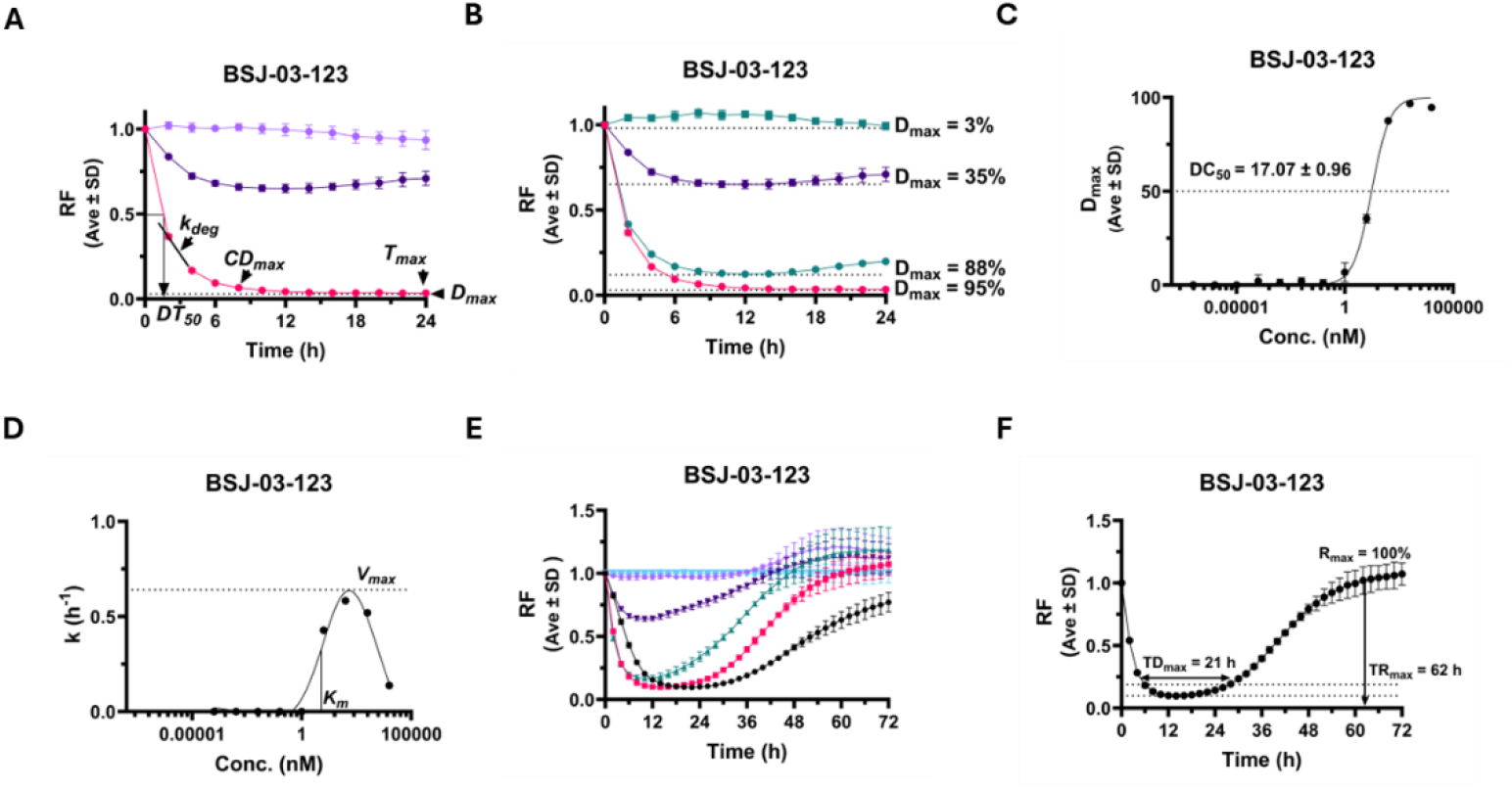
Degradation parameters A) Representative plot showing parameters calculated - maximum degradation (D_max_) ;concentration at maximum degradation (CD_max_); time at maximum degradation (T_max_); degradation rate (k_deg_) ; time at half degradation of the Protac (DT_50_) B) Plot showing D_max_ at different concentrations used to calculate DC_50_ C) Representative DC_50_ curve D) Representative graph showing maximum rate of degradation (V_max_) and half concentration needed to attain V_max_ (K_m_) E) Representative plot of dose-dependent effects observed over 72hours F) Representative plot showing time at D_max_ (TD_max_); maximum recovery (R_max_) and time to reach maximum recovery (TR_max_) obtained from a 72 hour study.

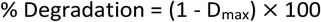

**DC**_**50**_ was calculated from a four-parameter fit of the D_max_ values at the various concentrations (Figs. 7B and S4A - I), plotted against their respective concentrations.^25^ A representative DC_50_ curve is shown in Fig. 7C and the curves for all PROTACs are shown in Figs. S5A – I. **k**_**deg**_ was obtained from the single component exponential decay fit of the degradation curve (at CD_max_ concentration) and was calculated as shown below

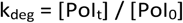

where [PoI_t_] is the concentration of the PoI at time t and [PoI_0_] is the concentration of the PoI at time zero. **DT**_**50**_ and ***τ*** were calculated from the k_deg_ as shown below

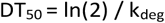

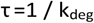

**V**_**max**_ and **K**_**m**_ were obtained by plotting k_deg_ as a function of concentration and fit using a non-linear regression analysis with a bell-shaped fit.^97^ A representative plot is shown in Fig. 7D and the curves for all PROTACs are shown in Figs. S6A - I. **TD**_**max**_, **R**_**max**_ and **TR**_**max**_ values were obtained from a 72-hour degradation curve (Figs. 7E - F and S7A – I, S8A - I). TD_max_ was calculated as shown below^78^

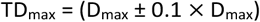

### Parameters obtained from Biophysical assays

The diversity observed in the dose-response degradation curves prompted us to explore the binding affinities (KdELECT) (Fig. S9A) and kinase activities (Kinase Hotspot) (Fig. S9B)^100^ of the PROTACs for CDK6. The data from these studies are summarized in Fig. S9C. The accumulation (Kp) of PROTACs in cells can be calculated from the binary binding constant as shown below^52^

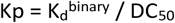

An AlphaLISA assay was done to assess the TC formation ability of the PROTACs (Fig. 8A). The fraction of the PROTAC bound in the TC (**θ**) was calculated as shown below

**Fig 8.**
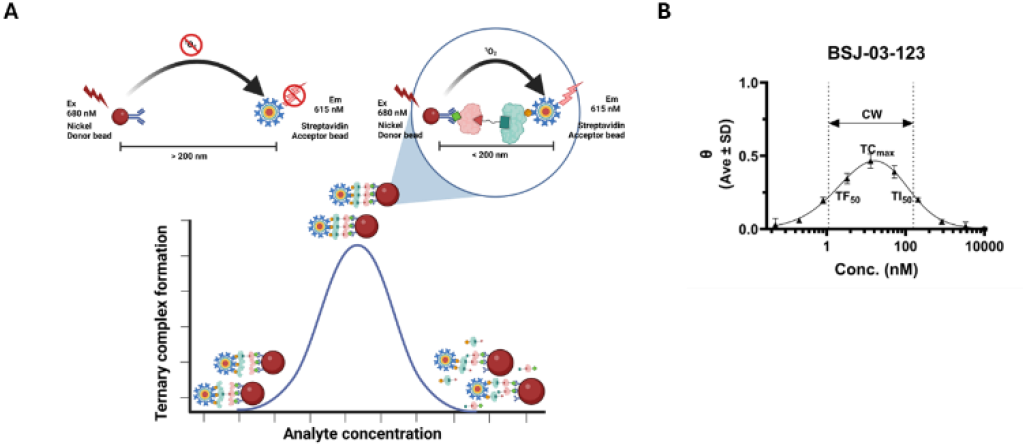
AlphaLISA A) Schematic of the assay B) Parameters calculated from an AlphaLISA TC assay – half maximum formation of TC (TF_50_); half maximum inhibition of TC (TI_50_); concentration at which maximum TC formation is seen (TC_max_) and the concentration window (CW) where TC formation is favorable.

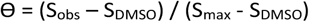

where the S_obs_ is the Alpha count for the PROTAC, S_DMSO_ is the Alpha count for DMSO and S_max_ is the maximum signal observed in the assay and S_max_ - S_DMSO_ is the assay window The Area under the Curve (**AUC**), was obtained by plotting the θ value for each PROTAC against concentration and fitting the curve using a non-linear regression, with a bell-shaped fit, consistent with three-body binding, on GraphPad Prism (Fig S10A). A four-parameter Sigmoidal fit of the rising portion of the bell curve was used to calculate the half-maximum concentration required for TC formation (**TF**_**50**_),^24^and a fit of the downward slope of the curve was used to calculate the half-maximum concentration required for TC inhibition (**TI**_**50**_) (Figs. 8B and S10C - G). The concentration at which maximum TC formation was observed (**TC**_**max**_) and the Concentration Window (**CW)** where effective TC formation was observed were obtained from the Alpha curve. CW was calculated as shown below

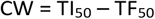

The values obtained from the ALPHA TC formation assay are summarized in Fig. S10H

## Discussion

Unlike traditional ligand binding curves that typically follow a Sigmoidal shape,^101^ PROTAC binding curves exhibit a bell shape (Fig. 5A).^102^ This biphasic response, described as the Hook effect,^24^ is the distinct difference between a PROTAC and an inhibitor and emphasizes the critical role of concentration selection for preliminary screens. The broad concentration range used for this screen captured responses from no degradation to almost complete degradation. As expected, all the CDK6 PROTACs showed a clear dose-dependent response with a Hook effect at higher concentrations, consistent with the Alpha assay results (Fig. S10A).

Variations in the initial degradation profiles reflect differences in either cell permeability, rate of TC formation and/or rate of primary ubiquitination.^78, 103^ However, variations at later time points could imply PROTAC instability (chemical or metabolic),^78^ reduced cellular PROTAC exposure potentially due to efflux,^104^ limited E3L recycling,^15^ differences in ubiquitination kinetics ^78^ or proteasomal capacity limitations.^105^

Notably, at the highest concentration, several (> 75%) PROTACs exhibited altered degradation profiles, probably due to concentration dependent kinetic limitations including reduced rate of TC formation due to the Hook effect, formation of non-cooperative complexes or “occupancy without degradation” stabilization of the TC.^45^ Though TC stability is essential for productive ubiquitination, excessive stability can limit PROTAC recycling and impair degradation.^106^ Therefore, an optimal balance is required, whereby the TC formed is sufficiently stable to allow the ubiquitination but can subsequently dissociate to allow for degradation.^5, 107^ Initial signal increases observed at higher concentrations, may reflect the stabilization of CDK6 within the TC, while increases above baseline observed at later time points may reflect an inhibitor-like effect of the PROTAC.^108^ These patterns are consistent with dose-dependent TC behavior, however, the study did not directly quantify TC occupancy in cells.

The diversity in the dose response degradation profiles highlights the complexity of the cellular mechanisms involved in PROTAC-mediated degradation and underscores the value of kinetic assays as functional readouts. Dynamic behaviors captured through a kinetic assay are often missed in static, single-read, or endpoint assays.

A typical PROTAC degradation curve exhibits three phases - an initial drop, followed by a plateau and eventually recovery, typically in experiments lasting beyond 24 hours.^24^ Good degraders often display the classic degradation profile (rapid, sustained and almost complete PoI depletion followed by slow PoI recovery upon PROTAC removal).^96^ PROTACs can therefore be classified using a combination of the following metrics – rapid or slow and complete or partial, based on their degradation profiles (Fig. 9A). All the CDK6 PROTACs tested exhibited classic degradation profiles, indicating PROTAC stability, low degrader efflux, efficient, repeated TC formation and effective ubiquitination over 24-hours.^28, 68^ Sustained degradation further implies that the rate of CDK6-GFP degradation exceeded its rate of synthesis.^48, 49^

**Figure 9.**
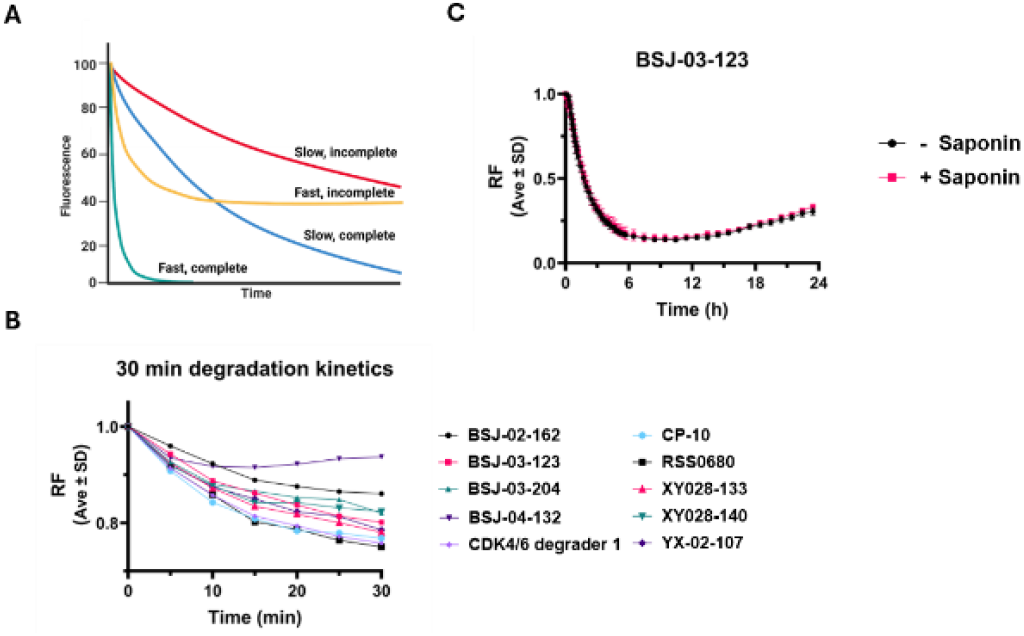
Classification plots A) Classification of degraders B) Initial degradation profile C) Permeability test.

Examination of the first 30 minutes of the degradation curve (Fig. 9B) showed that degradation onset occurred within 5 minutes, with appreciable loss (∼10%) of CDK6-GFP within 10 minutes. This suggests that the CDK6 PROTACs were cell-permeable, rapidly formed TCs, induced ubiquitination,^96^ and degradation. A permeability assay using Saponin pre-treatment showed no difference in the degradation profiles between treated and untreated samples (Figs. 9C and S11A - I), confirming the observed permeability. The observation that more than half of the CDK6 was degraded within 2 hours (Fig. S11J) by most (70%) of the PROTACs, confirms rapid PROTAC recycling.

Efficient degradation of CDK6-GFP (> 50%) was observed over a large concentration range (3 - 4 orders of magnitude) for most (75%) of the PROTACs, which is also reflected in their CW values, suggesting that these PROTACs could have a large therapeutic window. Conversely, a narrow CW would indicate that careful choice of concentration will be needed when screening such PROTACs.

BSJ-04-132 a CDK4 selective PROTAC, showed no TC formation at any of the tested concentrations, in the Alpha assay. However, 50% degradation of CDK6-GFP was observed in our degradation assay, likely due to the high sequence homology (∼70%) between CDK4 and CDK6.^109^ This elucidates how the complexity present in a cellular environment was not effectively captured in biophysical assays and that the cell based assay system minimized false negatives.

The ability to obtain indirect mechanistic insights, like permeability, formation, and stabilization of the TC and PROTAC recycling and therapeutic window from a single screen is an added benefit of this CDK6-GFP degradation assay.

### Efficacy

PROTACs can be ranked for efficacy based on their D_max_ value.^68^ While this type of ranking can be a useful cutoff during preliminary screens, ranking PROTACs with similar D_max_ values, particularly during hit-to-lead optimization, requires additional parameters.

Despite having similar D_max_ values, the rate of recovery and R_max_ values varied among the PROTACs evaluated, (Fig. 10A) indicating that R_max_ would be a better metric to rank hit PROTACs based on efficacy (Fig. 10B). VHL-based PROTACs exhibited a lower R_max_ (< 50%) indicating greater stability. A washout experiment (Figs. 10C and S12A - I) confirmed that sustained exposure to the PROTAC was necessary for degradation.^35,36,95^ Despite PROTAC removal from the media, continued, albeit limited CDK6-GFP degradation was observed for > 60% of the CDK6 PROTACs, resulting in incomplete CDK6-GFP recovery – consistent with their classic degrader profile. Together with the lack of change in the degradation profile following saponin treatment, these findings suggest that the washout kinetics likely reflect intracellular PROTAC exposure.

**Figure 10.**
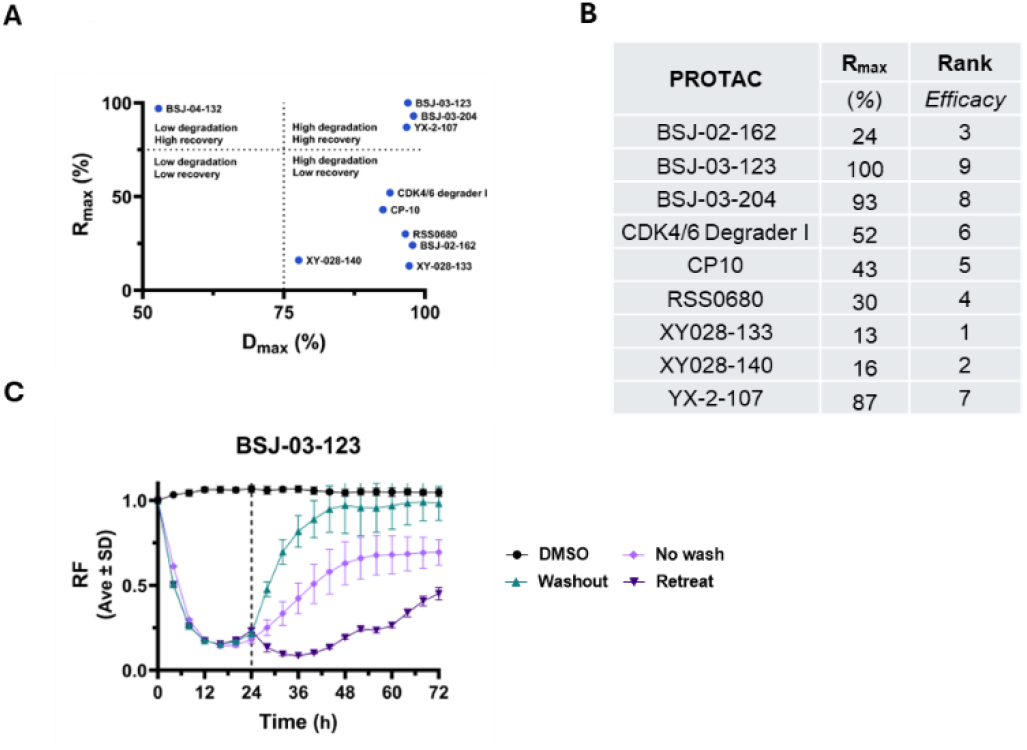
Efficacy A) Comparison of degradation (D_max_) and recovery (R_max_) B) Ranking based of efficacy (R_max_) C) Effect of washout (at 24 hours) and PROTAC retreatment.

Sustained degradation is a desirable quality for PROTACs. Recovery of PoI levels, following PROTAC treatment is typically observed as a consequence of declining intracellular PROTAC exposure due to compound instability (chemical or metabolic) ^110^ or efflux, limitations of the ubiquitin-proteasome system^108, 111^ or increased PoI synthesis. Most (>80 %) of the CDK6 PROTACs showed a TD_max_ greater than 24 hours and almost complete recovery of CDK6-GFP was observed with PROTACs having TD_max_ less than 24 hours. PROTACs having a carboxamide linker, greater than three oxygen atoms or a CRBN E3L showed decreased stability (TD_max_ < 24 hours) like previously reported.^112-114^

The relative stability, efflux, and intracellular exposure of PROTAC can be inferred from recovery and washout kinetics without direct quantification, representing another salient feature of the CDK6-GFP assay.

### Potency

DC_50_ is commonly used as a measure of PROTAC degradation potency. Despite having similar D_max_ values, the CDK6 PROTACs had different CD_max_ values, highlighting the importance of performing dose response preliminary screens to avoid missing hit compounds. The lack of correlation between the TC_max_ and CD_max_ values further emphasizes that caution should be exercised when making extrapolations from biophysical to cellular assays.

The rate of degradation of the PoI varies until it reaches a steady state at D_max_. Therefore, DC_50_ values calculated from the D_max_ values obtained for different concentrations is a better choice, because it is not time dependent (Figs. 7C and S5A - I).^25^ The DC_50_ trends observed for the CDK6 PROTACs were consistent with their CD_max_ trends, indicating that CD_max_ could also be a good indicator of potency.

Comparison of the imaging-derived DC_50_ against the 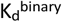 from the KdELECT assay showed good correlation (R^2^ - 0.79) (Fig. S13A), consistent with a first-pass association in early discovery. However, as the cellular DC_50_ integrates permeability, efflux, and ternary complex properties, 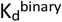 alone may not be determinative, as suggested by the weak TC_max_ - CD_max_ relationship. Consistent with the classic degrader profile, CDK6 PROTACs with higher potency showed lesser recovery (Fig. 11A) and PROTACs with a short linker length had a lower K_m_ value (Fig. S13B).

**Figure 11.**
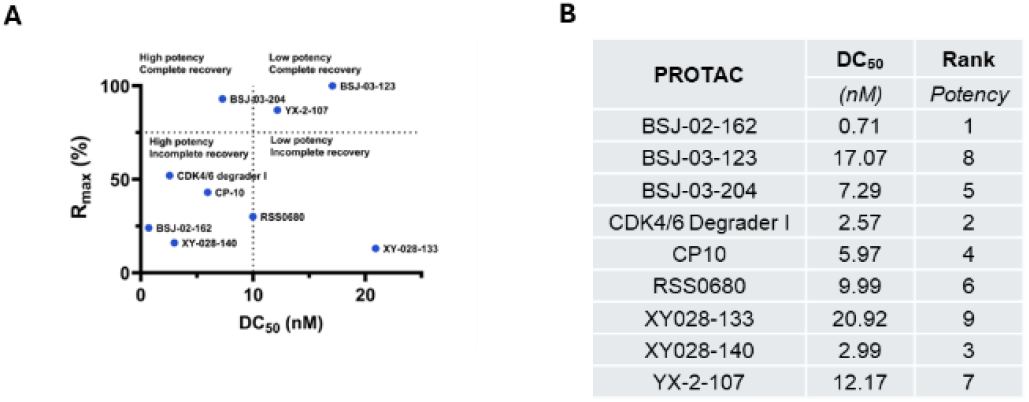
Potency – A) Comparison of efficacy (R_max_) and potency (DC_50_) B) Ranking based on potency (DC_50_).

Previous reports suggest that PROTACs can be considered catalytic if their cellular DC_50_ is less than their 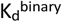.^115^ All the CDK6 PROTACs tested were catalytic, except YX-2-107, and the observed aberration could be explained by its low cellular accumulation. Conversely, high cellular accumulation could explain the excellent degradation observed for CP-10 and XY028-140 despite their moderate biophysical characteristics. Potency ranking of the CDK6 PROTACs based on their DC_50_ is shown in Fig. 11B.

### Efficiency

Although the rate of degradation of PROTACs is understood to play a key role in the degradation kinetics, efficiency is typically not considered when ranking PROTACs.

Despite significant protein loss (> 75%) within 6 hours, T_max_ was not observed until 22-24 hours for most of the PROTACs, implying that the degradation followed pseudo-first order decay kinetics.^78^ The observed DT_50_ of 2 hours indicated high degradation efficiency.^45^

The degradation of a PoI by a PROTAC can be approximated to enzyme catalysis if the affinity of the PROTAC for one binding partner is significantly greater than the other.^116^ In this case, the PROTAC bound CDK6 would function as the substrate for CRBN/VHL. Enzymatic decay of a substrate, described by the Michaelis Menten equation, generates a saturable Sigmoidal curve with increasing concentration of the substrate.^117^ However for PROTACs, because of the multiple equilibria, co-operativity and Hook effects, an adaptation of this model is needed. Recently a mathematical model was proposed to extend the Michaelis Menten concept to ternary systems like PROTACs.^97^ Most (> 60%) of the CDK6 PROTACs were fast, complete degraders of CDK6 (Fig. 12A). However, no correlation was observed between k_deg_ and R_max_ (Fig. S14A), DC_50_ (Fig. S14B), K_d_^binary^ or IC_50_, implying that the rate of degradation of CDK6-GFP (at CD_max_ concentration) was independent of the affinity, activity, or potency of the CDK6 PROTACs.

**Figure 12.**
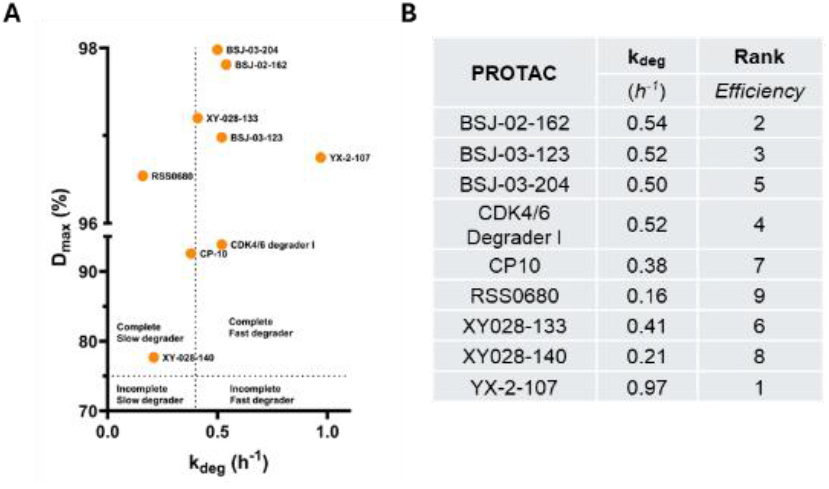
Efficiency – A) Comparison of degradation (D_max_) and efficiency (k_deg_) B) Ranking based on efficiency (k_deg_).

In drug pharmacology, the residence time (***τ***) is the amount of time a drug spends bound to the target.^118^ By extension, we can approximate *τ* for PROTAC degradation to be the time a PROTAC spends bound to the PoI, which can be used as a measure of the stability of the TC. A higher TC turnover (smaller *τ*) coupled with a higher TC formation (lower TF_50_) leads to increased PROTAC recycling and consequently, more efficient degradation (YX-2-107). Conversely, a larger *τ* and TF_50_ would indicate a slow degrader (XY028-140). Both conclusions were validated from the CDK6-GFP degradation profiles of the PROTACs. The efficiency ranking for the CDK6 PROTACs based on k_deg_ values is shown in Fig. 12B.

The CDK6 PROTACs ranked differently for efficacy (Fig. 10B), potency (Fig. 11B) and efficiency (Fig. 12B) highlighting the complex cellular degradation dynamics. Therefore, an equally weighted, comprehensive ranking which accounts for all three metrics would be a better method to rank PROTACs. All the calculated values obtained from the two screens and the overall ranking for the CDK6 PROTACs are shown in Fig. 13.

**Figure 13.**
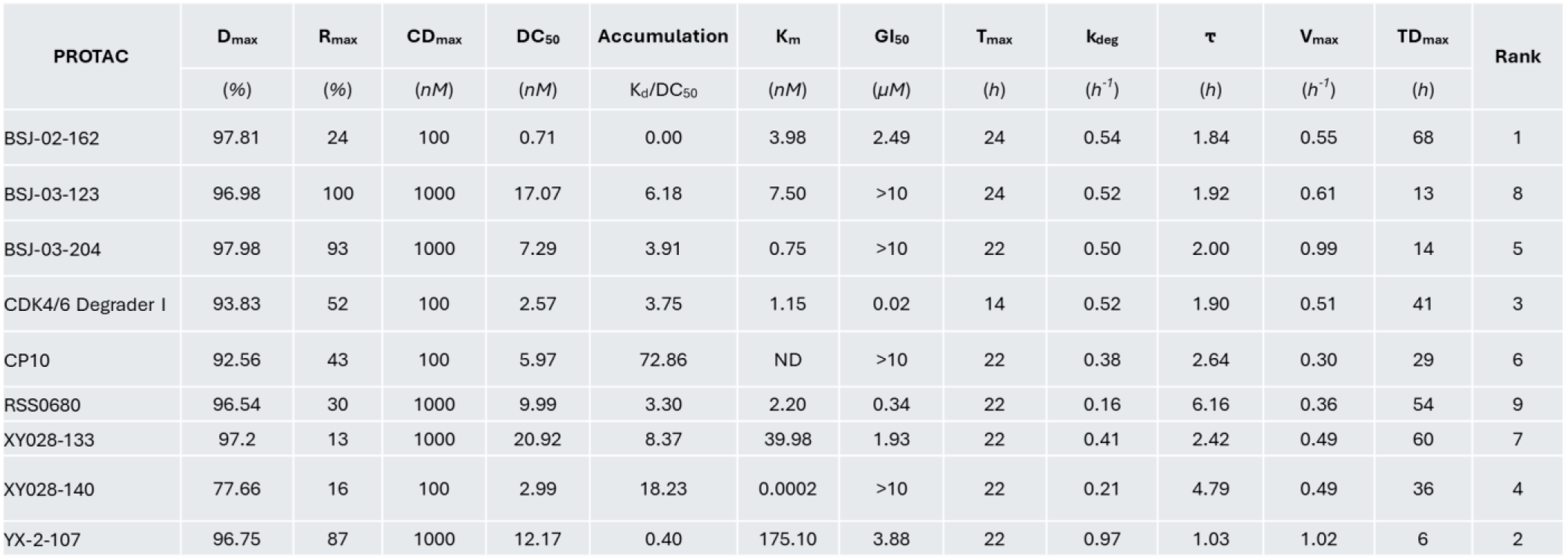
Table showing all the parameters obtained from the degradation assay and a comprehensive ranking of the PROTACs based on efficacy (R_max_), potency (DC_50_) and efficiency (k_deg_).

### Mechanistic assays

Additionally, the CDK6-GFP degradation assay was also able to detect competition (by either PoI or E3L ligand) and rescue (by either MLN4924^119^ or MG-132)^105^ (Fig. 14A) results.

**Figure 14.**
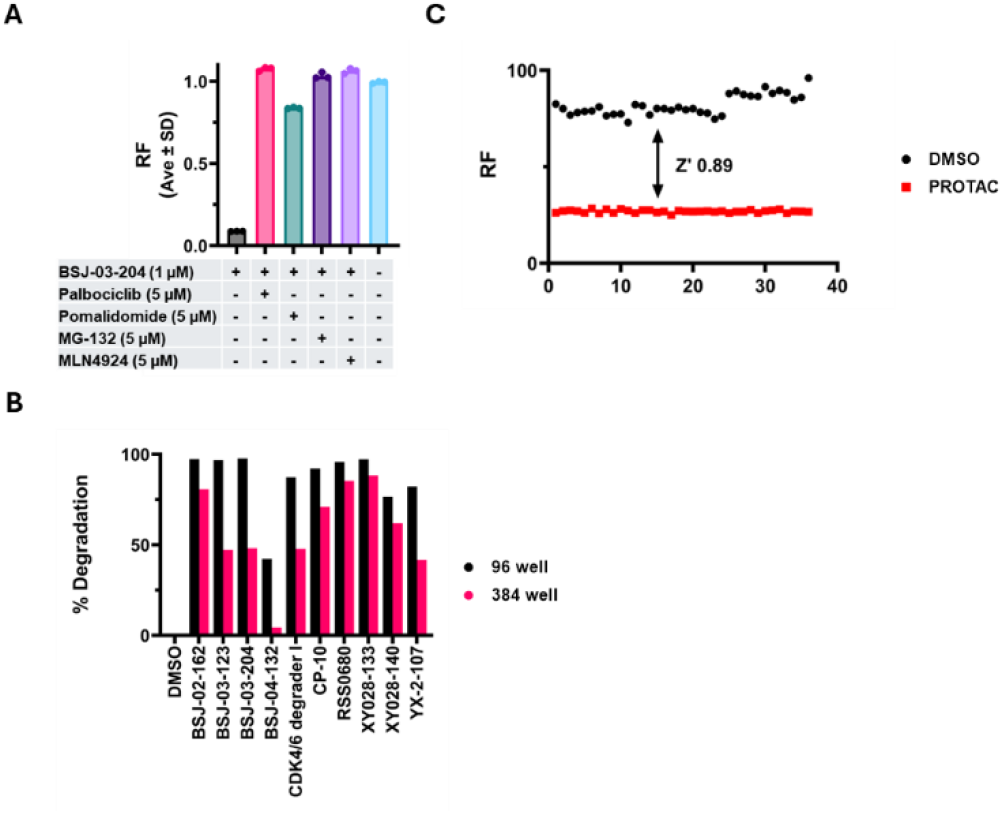
Mechanistic validation A) Competition and rescue with PoI, E3L ligand, MG-132 and MLN4924 B) Miniaturization C) HTS amenability

The ability to measure competition and complete rescue of CDK6-GFP implies that the MCF7 CDK6-GFP assay can be used to establish PROTAC mechanism of action (MoA).

### HTS compatibility

Comparable degradation trends between assays performed in 96-well and 384-well plates (Figs. 14B and S15A - I) indicate that the assay is robust and suitable for miniaturization. The primary assay validation metrics, and a Z’-factor of 0.84 (Fig. 14C), demonstrated that this assay was amenable to HTS.^33^

### Orthogonal assays

Orthogonal assays were performed to compare and correlate the degradation of CDK6 seen in the live-cell imaging assay with other available techniques.

As fluorescence was the key readout of the CDK6-GFP imaging assay, we measured the GFP fluorescence of cell lysates obtained after a 1 µM, 24-hour PROTAC treatment on a plate reader (Fig. 15A). The percent degradation obtained (Fig. S16B) corelated well the imaging experiments (Fig. S16A) (R^2^ – 0.97) indicating that the MCF7-CDK6-GFP reporter system is also amenable to simple non-imaging fluorescent detection techniques.

**Figure 15.**
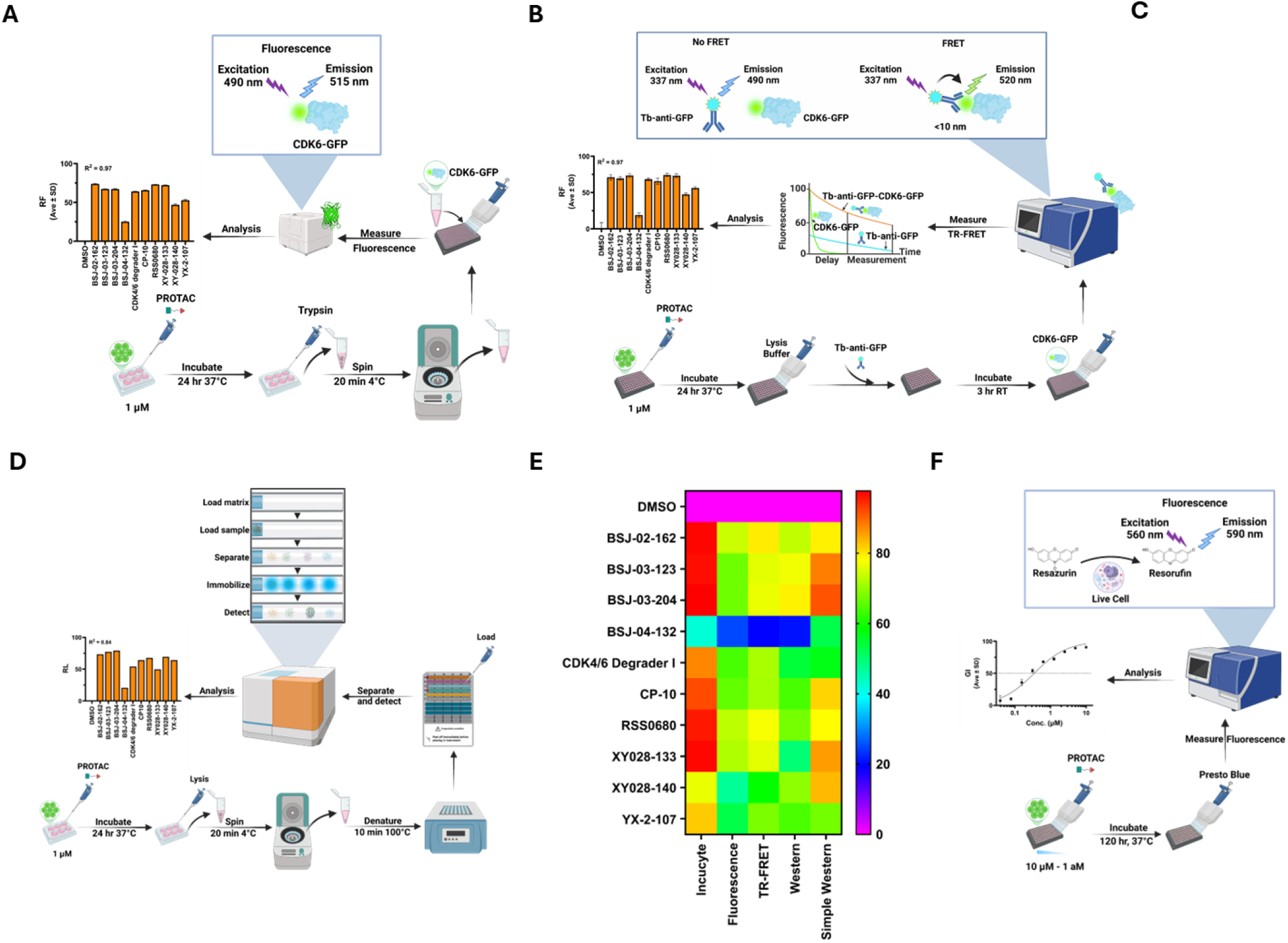
Orthogonal comparative assays with 1 µM 24-hour treatment of PROTACs A) Fluorescence B) TR-FRET Lanthascreen C) Traditional Western with MCF7-CDK6-GFP D) Simple Western with MCF7-CDK6-GFP or BxPC3 E) Percentage degradation across all tested assays F) Growth Inhibition

Proximity based assays are valuable tools used to study PROTAC interactions. A TR-FRET assay was performed on lysates obtained after a 1 µM, 24-hour PROTAC treatment, lysed in the presence of a Tb-GFP antibody. The energy transfer from the Tb donor to the GFP acceptor occurs only if the two are in proximity (< 10 nm) (Fig. 15B). The time-resolved fluorescence emitted was measured and the FRET ratio was calculated as shown below

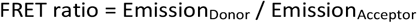

The percentage degradation obtained from the TR-FRET assay (Fig. S16C), corelated well (R^2^ – 0.96) with the imaging experiments (Fig. S16B).

After confirming that the GFP loss seen in the live-cell imaging was measurable with two other fluorescent techniques, we proceeded to ensure that loss of CDK6 could be measured by non-fluorescent methods. First, a traditional immunoblot was performed by probing lysates obtained after a 1 µM, 24-hour PROTAC treatment for CDK6 (Fig. 15C). The trend for degradation was present (Fig. S16D) with reasonable correlation (R^2^ – 0.77). Next, an automated capillary electrophoresis was performed with the same treatment (Fig. 15D) and the degradation seen (Fig. S16E) showed excellent correlation (R^2^ – 0.84) with the imaging assay (Fig. S16A). Overall, the automated method showed better correlation to the CDK6-GFP degradation assay results.

Having established that both GFP and CDK6 could be detected in non-imaging settings, we wanted to confirm the degradation of CDK6 by the PROTACs in a non-fused endogenous system. We used a Pancreatic Cancer cell line (BxPC3) expressing high endogenous levels of CDK6. The PROTACs performed predictably (Fig. S16F) with excellent correlation (R^2^ – 0.86) confirming that the loss of GFP signal seen in the imaging assay reflected CDK6 loss.

All orthogonal assays done showed strong agreement (Fig. 15E) supporting the robustness and translatability across techniques under evaluated conditions.^54^ The rankings of the PROTACs based on the orthogonal assays are shown in Fig. S16G.

Finally, a growth inhibition assay was performed over a 5-day period to assess if the PROTAC mediated CDK6 degradation inhibited cell growth. Cells were treated with a 2-fold 9-point dilution series of each PROTAC (10^-6^ to 10^-9^ M). Cell viability was measured at the beginning of PROTAC addition (T_0_) and after 120 hours (T_100_). The percentage growth and percentage growth inhibition (GI_50_) were calculated as shown below.

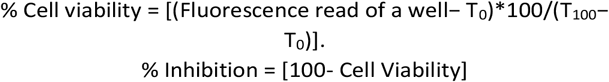

The percent inhibition was plotted as a function of the concentration and fit with a four-parameter Sigmoidal fit on GraphPad Prism and the GI_50_ was obtained.

The results showed that half the PROTACs assessed had no effect on growth inhibition. The presence of either Thalidomide or VHL ligands conferred low micromolar growth inhibitory effects, and the trends reported,^51, 88, 89, 94^ were comparable to what was obtained in MCF7-CDK6-GFP cells (Fig 14G). However, CP-10,^90^ XY028-140^93^ and XY028-133^60^ showed contrary results to what was reported. The GI_50_ values are summarized in Fig. S16H.ARTICLE

## Conclusions

The FDA-approval of the first PROTAC earlier this year ushers in a new era for PROTACs, bringing with it the need for better tools for hit-finding, hit-to-lead optimization and MoA studies.

Here, using a panel of commercially available CDK6 degraders, we demonstrated how an MCF7 cell line stably expressing CDK6-GFP can be used to rapidly generate dynamic degradation metrics (D_max_, R_max_, TD_max_, CD_max_, DC_50_, K_m_, T_max_, K_deg_, V_max_, DT_50_, *τ* and TR_max_) for comparing and ranking PROTACs within a controlled reporter system. The observed CDK6-GFP degradation was orthogonally validated in an endogenous non-fused system.

We also demonstrated that the PROTACs exhibited distinct degradation signatures and that kinetic dose response analysis enabled the determination of key parameters (R_max_, DC_50_ and k_deg_). These parameters were essential to accurately rank the PROTACs, based on their individual metrics of efficacy, potency, and efficiency, as well as comprehensively. This comprehensive ranking accounts for the different modalities of action and therefore provides a better metric to rank PROTACs both for hit-finding and hit-to-lead optimization.

Other parameters like permeability, TC formation and stabilization, PROTAC stability, recycling, cellular exposure, efflux, and therapeutic window could be inferred from the degradation profiles. TC-related behavior was supported indirectly by alignment with experimental AlphaLISA and competition/rescue but was not directly measured in cells.

This comprehensive study of CDK6 PROTACs showed that R_max_ is a better indicator of PROTAC efficacy; VHL PROTACs with fewer oxygens and amine linkers are preferred for long term suppression of CDK6, while CRBN PROTACs with a carboxamide linker are suitable for transient CDK6 degradation; thalidomide PROTACs were more potent, and PEG linkers suffer from stability issues.

The ease of use, rapid assay turnaround time coupled with the lack of additional reagents needed are the most attractive aspects of this assay. Additionally, the robustness and translatability of the MCF7 CDK6-GFP degradation to non-imaging techniques will support wider applicability and the calculated kinetic parameters associated with the degradation process provide insights that may be extended to other systems.

While a considerable limitation is that the GFP binding site on the PoI could interfere with the native function of the protein, requiring alternate systems to study functional consequence of the PoI degradation; this report provides compelling evidence to expand the use of GFP fused PoI based assay systems for discovery and development of PROTACs.

### Experimental

Details of the materials used are provided in *Supplementary Table 1*.

#### Cell culture

MCF7 WT and MCF7-CDK6-GFP cells were cultured in DMEM-High containing 10% Fetal Bovine Serum (FBS), 1% Penicillin-Streptomycin (Pen-Strep) and human Insulin (10 µg/mL). MCF7-CDK6-GFP cells were supplemented with Puromycin (2 µg/mL). BxPC-3 was cultured in RPMI containing 10% FBS and 1% Pen-Strep. Cell lines were cultured at 37°C, in a humidified incubator in the presence of 5% CO_2_. Cells were routinely evaluated for Mycoplasma using the MycoAlert mycoplasma detection kit. For imaging studies, cells were plated in DMEM Fluorobrite containing 10% FBS and 1% Pen-Strep.

#### Generation of CDK6-GFP cell lines

10,000 MCF7 cells/well were plated in 200 µL media in a sterile, clear 48-well plate and allowed to adhere overnight. The next day, 8 µg/mL polybrene was added to the media before virus inoculation. Human CDK6-GFP lentiviral particles were thawed on ice and gently mixed. Varying ranges of MOI (10 - 1.25) of the virus were added to the MCF7 cells and gently mixed. For reverse transduction, the same range of virus concentration was used on freshly trypsinized cell suspensions. The plate containing the cells and virus was spun in a centrifuge for 30 min at 1000g at room temperature and then transferred to an incubator for 72 hours. The media was replaced after 72 hours, with media containing 2 µg/mL puromycin. The cells were selected with puromycin for 2 weeks before screening. The presence of GFP was confirmed by visualizing on an Evos™ M5000 microscope (Thermo Fisher, USA). Images to compare morphology were taken at 10X magnification on a Zeiss Axiovert 40C microscope (Zeiss, Germany).

#### Screening for growth variations

30,000 cells/well were plated in 100 µL growth media in a 96-well sterile, black-walled, clear-bottom plate and allowed to adhere overnight. The next day, they were screened for growth (Phase channel) and CDK6-GFP expression (Green channel) on an Incucyte^®^ SX3 (Sartorius Germany). Whole well scans at 4X magnification were done every 2 hours for 24 - 72 hours. Masks were applied, and any cell showing Green calibration Unit (GCU) greater than 0.5 was considered a signal. The background GCU was typically 0. The data was exported to Microsoft Excel. The fluorescence data was normalized to the confluence to correct for growth differences. The data was further normalized to DMSO control at each time point to remove any DMSO contribution to the effect seen. Further normalization to time zero accounted for any seeding irregularities. The normalized data was then plotted on GraphPad Prism (**version 10.6**) and percentage degradation was calculated.

#### Flow Cytometry

1,000,000 cells were trypsinized, pelleted (5 min at 200 g), and rinsed once with 1X Phosphate Buffered Saline (PBS). The supernatant was removed by aspiration, and 500 µl of cold PBS was added and mixed gently. A modified protocol for GFP containing cells was used to process samples.^120^ 500 µL of ice-cold 2% paraformaldehyde was added and mixed again. The cell suspension was incubated for an hour at 4°C. The cells were then centrifuged (5 min at 300 g, at 4°C) and the supernatant was removed by aspiration. The pellet was washed once with cold 3 mL PBS. The cells were then permeabilized by the dropwise addition of 1 mL of 70% ethanol, while gently vortexing. Care was taken to ensure single cell suspensions were obtained by ensuring continuous vortexing while adding ethanol. The cell suspension was incubated overnight at 4°C. The cells were then centrifuged (5 min at 300 g, at 4°C) and the supernatant was removed by aspiration. The pellet was washed once with cold 3 mL PBS. 1 mL of propidium iodide diluted in PBS (final concentration 40 µg/mL) was added, and the cell suspension was incubated in the dark for 30 min at 37°C. Samples were filtered through a 35 µm nylon mesh to remove clumps and immediately analyzed on the FACSCalibur (BD Biosciences). The data obtained was processed on ModFit LTTM (Verity Software House) and plotted on GraphPad Prism.

#### Western Blot

1,000,000 cells were trypsinized, pelleted and rinsed with 1X PBS. The pellets were lysed by adding 75 µL of RIPA buffer (containing protease and phosphatase inhibitors) and gently breaking down the pellet completely. The suspension was incubated on ice for 30 min with intermittent vortexing for 2 seconds every 10 minutes. Lysates were spun down at 4°C for 20 minutes at 17000 rpm. The clear supernatant was transferred to a clean, chilled tube. Protein quantitation was done using the BCA method. 25-40 µg of protein was loaded per well onto a 4 - 15% gel and separated at 120V for 90 min. A kaleidoscope ladder was run to estimate the molecular weights of the separated proteins. The gel was washed at 4°C with water for 5 min and then with cold transfer buffer for 10 min. The separated proteins were transferred to a PVDF membrane for 8 min and blocked for an hour at room temperature in 5% non-fat milk or 5% Bovine Serum Albumin in 1X Tris Buffered Saline with 0.01% Tween (TBST). The membrane was incubated in the primary antibody for 16 hours at 4°C and then washed for 30 minutes (3X for 10 minutes each) with 1X TBST. The membrane was then incubated with the respective secondary antibody for 1 hour at room temperature. Afterward, the membrane was washed again for another 30 minutes (3X for 10 minutes each) with 1X TBST and then probed with ECL and imaged on a ChemiDoc MP (Bio-Rad, CA). A list of the antibodies used, and their dilutions are provided in *Supplementary Table 2*. The images were quantified using ImageJ (NIH). The correlation plots plotted on GraphPad Prism.

#### Simple Western

Cells lysates were prepared as stated in the Western Blot assay section. 0.1X sample buffer was prepared by diluting the provided 10X sample buffer with distilled water. The lysates were diluted with 0.1X sample buffer to a final concentration of 1 mg/mL. 5X Fluorescent Master Mix (FMM) was prepared by adding a 1:1 mixture of 10X sample buffer and 400mM dithiothreitol (DTT). FMM was spiked into each sample to a final concentration of 1X. The samples were then heated on a heating block at 100°C for 10 minutes and then chilled on ice for 10 minutes. The samples were briefly spun down and 6 µL of each sample was loaded per well of the 384-well plate. The Biotin ladder was loaded in the first lane. The remainder of the plate was filled per the manufacturer’s instructions (https://www.bio-techne.com/pdf-download-arena-document/product-insert/pl3-0005) A 12-230 kDa separation module for size separations was used to separate and mobilize the proteins on a Jess (Bio-techne, MN). Suitable dilutions of the primary antibody were made in Antibody diluent (following optimization) and used with the appropriate secondary antibodies provided by the manufacturer. Secondary antibodies were used “as is” without further dilution. A complete list of the antibodies used, and their dilutions are provided in supplementary table 2. The separated bands were visualized using a 1:1 mix of Luminol-S and Peroxide. Loading controls were run using the Replex module reagents per manufacturer’s instructions. The data obtained was portrayed as lane images using the Compass for Simple Western software (version 5.0.1). The Area under the curve (AUC) was used as a measure of the band intensity. The sample bands were first normalized to the loading control to account for any loading variations. Subsequently, they were normalized to the DMSO sample to calculate the fold-change in the expression level of each protein. The percent degradation (for degradation assays) was calculated as previously shown, and correlation plots were plotted on GraphPad Prism.

#### Assay validation

30,000 cells/well were plated in 90 µL Fluorobrite media in 96-well, black-walled plates. To determine assay noise, whole well scans of 60 wells (phase and green channel) were done, at 4X magnification, every 4 hours for 72 hours. Images were collected, and the data was processed as previously stated in the preliminary screening.

The linearity experiment was done by half-diluting 40,000 cells/well in 100 µL Fluorobrite media in in 96-well, black-walled plates. The experiment was performed in triplicate and cell free blank wells, containing only media (100 µL), were also included. The plates were imaged, and the data was processed as previously stated.

30,000 cells/well were plated in 90 µL Fluorobrite media in 96-well, black-walled plates for the DMSO tolerance experiment. They were treated, in triplicate, with half-dilutions of DMSO starting at 10% and DMSO free wells were included as controls. Whole well scans (phase and green channel) were done, at 4X magnification, every 4 hours for 72 hours. Images were collected, and the data was processed as previously described.

To measure precision, inter and intraday variation, 30,000 cells/well were plated in 90 µL Fluorobrite media. 40 wells were treated with 0.1% DMSO and imaged (phase and green channel), and the data was processed as previously stated.

#### Degrader screening

10 mM stock solutions of the degraders were prepared in DMSO and aliquoted to prevent multiple freeze-thaws. 25 - 30,000 cells/well were plated in 90 µL Fluorobrite media. The next day, PROTAC dilutions were made in DMSO before mixing with warm media to give 10X the final desired concentration. 10 µL of the media-diluted 10X PROTACs were added to the cells, in triplicate, to give a final DMSO concentration of 0.1%. DMSO controls were also done for all runs. Whole well scans at 4X magnification were done every 2 hours for 24 - 72 hours. Data was processed as previously stated.

#### Binding affinity

The biding affinity 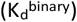 of CDK6 was calculated from an 11 point 3-fold compound dilution using the KdELECT™ competition binding assay performed by Eurofins. Binding constants 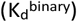 were calculated with a standard dose-response curve using the Hill equation

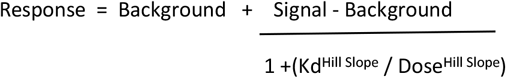

The Hill Slope was set to -1 and curves were fitted using a non-linear least square fit with the Levenberg-Marquardt algorithm.

#### Kinase activity

A 10 point 5-fold PROTAC dilution was done in triplicate starting at 10 µM at K_m_ concentrations of ATP. Staurosporine was used as the control compound and the data obtained was normalized to DMSO and curve fits were performed where the enzyme activity at the highest PROTAC concentration was less than 65% and the IC_50_ was calculated.

#### AlphaLISA

The CDK6-CRBN PROTAC optimization kit was used to perform this assay in 384-shallow well ProxiPlates with a final DMSO concentration of 0.1%. 2.5 µL of an 8 point 3-fold dilution was made for each PROTAC, in duplicate, in the Lysis buffer provided by the manufacturer, at 4X the final desired concentration (10 µM – 50 nM). DMSO controls were was also done for all runs. 2.5 µL of a master mix containing 240 nM each of His-CDK6/CyclinD3 and CRBN-Biotin (final concentration of each protein in the reaction was 30 nM) was added to each well and the three-component mixture was co-incubated for 1 hour at room temperature with gentle mixing (100 rpm orbital mixing for 30 seconds every 5 minutes). 2.5 µL (20 µg/mL final concentration) of Nickel Donor beads were added to each well and incubated in the dark, at room temperature for 60 minutes with mild occasional agitation (100 rpm orbital mixing for 30 seconds every 15 min). 2.5 µL (20 µg/mL final concentration) of Streptavidin Acceptor beads were then added to each well and incubated in the dark, at room temperature for 60 minutes with mild agitation (100 rpm orbital mixing for 30 seconds every 15 min). Care was taken to perform all parts of the experiment under low lighting conditions. Alpha counts were measured on a CLARIOstar Plus (BMG Labtech, NC) (Ex 680 nm for 180 ms, 0.02 ms delay, 634 nm dichroic, Em 615 nm with 20 ms integration). The data was plotted against the concentration to get curves using GraphPad Prism and a 1/Y2 weighting was performed per manufacturer’s instructions.

#### Permeability

30,000 cells/well were plated in 80 µL Fluorobrite media in sterile, black-walled, clear-bottom plates and allowed to adhere overnight in a humidified 5% CO_2_ incubator at 37°C. A 0.1% Saponin solution was made in sterile water and 10 µL of this solution was added to cells in triplicate. 10 µL of the media-diluted PROTACs at 10X CD_max_ concentration for each PROTAC was added to the cells, in triplicate. Phase channel and Green channel images were captured at 10X magnification every 15 minutes for 24 hours. Data was processed as previously discussed.

#### Washout

30,000 cells/well were plated and treated with the CD_max_ concentration of each PROTAC as outlined in the previous section. Images were collected every 2 hours, for 24 hours and the media was removed after 24 hours and replaced with fresh, warm media without PROTACs. The plates were returned to the incubator, and images were collected every hour for an additional 48 hours. Data was processed as stated before.

#### Competition and rescue

30,000 cells/well were plated in 90 µL Fluorobrite media. 10 mM stock solutions of Palbociclib, Ribociclib, Pomalidomide, Thalidomide, MG-132 and MLN-4924 were made in DMSO and aliquoted to prevent multiple freeze-thaws. The next day, 10X CDmax concentration of each PROTAC in DMSO was mixed with a dose dilution of each competitor in DMSO and warm media was added to give 10X the final concentration needed. 10 µL of the media-diluted PROTAC-Competitor mix was added to the cells in triplicate to give a final DMSO concentration of 0.2%. 0.2% DMSO controls were also done for all runs. The plates were then imaged and processed as stated before.

#### Miniaturization

20,000 cells/well were plated in 45 µL Fluorobrite media in sterile, black-walled, clear-bottom 384 well plates. PROTAC dilutions were made as previously described and 5 µL of the media-diluted PROTACs were added in triplicate to give a final DMSO concentration of 0.1%. DMSO controls were also done for all runs. The plates were imaged and the data collected was processed as previously stated.

#### Secondary validation

30,000 cells/well were plated in 90 µL Fluorobrite media. The cells were treated with 0.1% DMSO or 1 µM XY028-140. The plates were imaged (phase and green channel) every 4 hours for 72 hours and the data was collected and processed as previously described.

#### Fluorescence

1,000,000 cells/well were plated in 2 mL growth media in 6-well plates and treated with 1µM of each PROTACs. 24 hours later the cells were harvested and processed like a Western Blot. A 10-point 2-fold protein dilution curve was performed by half diluting the protein in lysis buffer. Lysis buffer was used as the negative control. Fluorescence (Ex 482, Em 520) was measured on a CLARIOstar Plus (BMG Labtech, NC). The data was plotted on GraphPad Prism to determine the concentration required for subsequent assays. 15 µL of a 6 mg/mL concentration of each lysate was prepared, in duplicate, by diluting the lysates in lysis buffer (if necessary). The fluorescence value of each sample was measured, and the data was normalized to the DMSO control to calculate the fold change. The percent degradation was calculated as previously shown, and correlation plots were plotted on GraphPad Prism.

#### TR-FRET

30,000 cells/well were plated in black-walled plates and treated with the CD_max_ concentration of each PROTAC as outlined in the previous section. After 24 hours of PROTAC treatment, the media was removed and 40 µL of complete HTRF lysis buffer was added to each well. Complete HTRF lysis buffer was prepared by adding 1X Protease and Phosphatase inhibitors to the Lysis buffer. The covered plate was incubated in the dark with agitation (300 rpm orbital mixing) for 30 minutes. 10 µL of complete lysis buffer containing 2 nM (final concentration) of Tb-anti-GFP antibody was added and gently mixed (100 rpm orbital mixing) for 30 seconds. The plate was stored, covered in the dark at room temperature for 3 hours. 40 µL of the lysate was then transferred to 384-well white opaque plates and fluorescence was measured on a Spectramax i3X outfitted with a TRF module. Acquisition conditions for the TR-FRET were obtained from the Tb-anti-GFP manufacturer’s site. The TR-FRET ratio of 520/490 (Acceptor emission/Donor Emission) was normalized to DMSO to get a fold change. The percent degradation was calculated, as previously stated, and plotted on GraphPad Prism to obtain correlation plots.

#### Growth Assay

3,000 cells/well were plated in 90 µL growth media in sterile, clear, flat-bottomed, 96-well tissue culture plates. The next day, PROTAC dilutions were made as stated in the degrader screening section and 10 µL of the media-diluted PROTACs were added to the cells in triplicate. 10 µL Presto blue was added per well to three control wells containing 0.1% DMSO alone and incubated in a humidified 5% CO_2_ incubator at 37°C for 15 min. Fluorescence (Ex 560, Em 590) was measured on a Spectramax i3X (Molecular Devices, CA) to determine T_0_. The plates were returned to the incubator for 72 hours. Post 72 hours, 10 µL Presto blue/well was added to all the remaining wells and incubated in a humidified 5% CO_2_ incubator at 37°C for 15 min. Fluorescence was measured on the i3X as before. Control wells with 0.1% DMSO alone were used to determine T_100_. The data obtained was plotted on GraphPad Prism, and the IC_50_ values were determined using a four-parameter Logistic curve-fitting function.

## Author contributions

Conceptualization, visualization formal analysis, editing draft and project administration: SK, AN; data curation, validation, investigation, methodology, writing original draft: SK; funding acquisition, resources, and review: AN

## Conflicts of interest

There are no conflicts to declare.

## Data availability

The data supporting this article has been included as part of the Supplementary Information and Appendix. Supplementary information: Materials used are in Tables S1 - 2 and supporting data is in Figures S1A – S16H. Raw western blots are in Appendix A - E

## Acknowledgements

The work done was supported in part by NIH R01CA197999.

## Notes

### Competing Interest Statement

The authors have declared no competing interest.

