## Supplementary Tables and figures for "Beyond D_max_ and DC_50_: A simple tool to evaluate the complex PROTAC-mediated kinetic degradation dynamics"

#### Supplementary materials

[a] *Division of Pharmaceutical Sciences, College of Pharmacy, University of Cincinnati, OH, 45267, USA*
[b] *Eppley Institute for Cancer Research, University of Nebraska Medical Center, Omaha NE, 68198, USA*

#### Table 1 - List of supplies

| S. No. | Item | Catalog No. | Company |
|---|---|---|---|
| 1 | MCF7 WT | **HTB-22** | ATCC |
| 2 | HEK293 | **CRL-1573** | ATCC |
| 3 | BxPC-3 | **CRL-1687** | ATCC |
| 4 | Human CDK6-GFP lentivirus | **RC208560L4V** | Origene |
| 5 | Tb-anti-GFP antibody | **A13391** | Invitrogen |
| 6 | DMEM-High | **SH30022.FS** | Cytiva |
| 7 | RPMI | **SH30027.FS** | Cytiva |
| 8 | Fluorobrite | **A1896701** | Thermo Fisher |
| 9 | OptiMEM | **31985062** | Gibco |
| 10 | FBS | **45001-106** | Corning |
| 11 | Pen-Strep | **16777-164** | Hyclone |
| 12 | Trypsin | **SH30042.01** | Cytiva |
| 13 | PBS | **SH30256.FS** | Cytiva |
| 14 | RIPA | **89900** | Pierce |
| 15 | Transfer buffer | **J63664.K3** | Thermo Fisher |
| 16 | TBS | **T60075-5000.0** | RPI Corp |
| 17 | Sample buffer | **042-195** | ProteinSimple |
| 18 | HTRF lysis buffer | **A12891** | Invitrogen |
| 19 | Insulin | **12585014** | Gibco |
| 20 | Puromycin | **A1113802** | Gibco |
| 21 | Hygromycin | **10687010** | Gibco |
| 22 | Polybrene | **TR-1003** | Sigma |
| 23 | Paraformaldehyde | **8187150100** | Sigma |
| 24 | 70% ethanol | **2701** | Decon Labs |
| 25 | Propidium iodide | **P3566** | Thermo Fisher |
| 26 | Protease and Phosphatase inhibitor (WB) | **78447** | Thermo Fisher |
| 27 | Non-fat milk | **MSPP-M0841** | Lab Scientific |
| 28 | Bovine Serum Albumin | **BP9706-100** | Fisher Scientific |
| 29 | Tween | **AAJ20605AP** | Fisher Scientific |
| 30 | Fluorescent Master Mix | **PS-FL01-8** | ProteinSimple |
| 31 | DTT | **PS-ST01EZ-8** | ProteinSimple |
| 32 | Antibody diluent | **042-203** | ProteinSimple |
| 33 | Luminol-S | **043-311** | ProteinSimple |
| 34 | Peroxide | **043-379** | ProteinSimple |
| 35 | Replex | **RP-001** | ProteinSimple |
| 36 | DMSO | **BP231-100** | Fisher Scientific |
| 37 | Protease inhibitor (TR-FRET) | **P8340** | Sigma |
| 38 | Phosphatase inhibitor (TR-FRET) | **P0044** | Sigma |
| 39 | MycoAlert kit | **LT07-318** | Lonza |
| 40 | BCA kit | **23227** | Pierce |
| 41 | ECL | **RPN2232** | Amersham |
| 42 | CDK6-CRBN PROTAC optimization kit | **79924** | BPS Biosciences |
| 43 | Nano-Glo Luciferase Assay System | **N1110** | Promega |
| 44 | Nylon mesh | **352235** | Falcon |
| 45 | PVDF membrane | **IPVH00010** | Millipore |
| 46 | 4 - 15% gel | **4561086** | Bio-Rad |
| 47 | Kaleidoscope ladder | **1610375** | Bio-Rad |
| 48 | Biotin ladder | **part of EZ Standard pack 1** | Protein Simple |

| | | | |
|---|---|---|---|
| 49 | 12-230 kDa separation module | **PS-CC01** | ProteinSimple |
| 50 | Nickel Donor beads | **AS101D** | Revvity |
| 51 | Streptavidin Acceptor beads | **AL125C** | Revvity |
| 52 | Presto blue | **A13262** | Invitrogen |
| 53 | Saponin | **SAE0073** | Sigma |
| 54 | BSJ-02-162 | **HY-144995** | MedChemExpress |
| 55 | BSJ-03-123 | **HY-111556** | MedChemExpress |
| 56 | BSJ-03-204 | **HY-136250** | MedChemExpress |
| 57 | BSJ-04-132 | **HY-136252** | MedChemExpress |
| 58 | CDK4/6degrader I | **HY-163786** | MedChemExpress |
| 59 | CP-10 | **HY-125835** | MedChemExpress |
| 60 | RSS0680 | **HY-148062** | MedChemExpress |
| 61 | XY028-133 | **HY-129180** | MedChemExpress |
| 62 | XY02-140 | **S9880** | Selleckchem |
| 63 | YX-2-107 | **HY-148530** | MedChemExpress |
| 64 | Palbociclib | **HY-50767** | MedChem Express |
| 65 | Ribociclib | **HY-15777** | MedChem Express |
| 66 | Pomalidomide | **HY-10984** | MedChem Express |
| 67 | Thalidomide | **HY-14658** | MedChem Express |
| 68 | MG-132 | **HY-13259** | MedChem Express |
| 69 | MLN-4924 | **HY-70062** | MedChem Express |
| 70 | 48-well sterile, clear plate | **150687** | Fisher Scientific |
| 71 | 96-well sterile, black-walled, clear-bottom plate | **3882** | Corning |
| 72 | 384-well plate for Jess | **PS-PP03** | Protein Simple |
| 73 | 384-shallow well ProxiPlates | **6008280** | Revvity |
| 74 | 96-well sterile, clear, flat-bottom plate | **12-565-438** | Fisher Scientific |
| 75 | 6-well sterile, clear, flat-bottom plate | **FB012927** | Fisher Scientific |
| 76 | 384-well white opaque plates | **6007290** | Revvity |
| 77 | 96-well sterile, white-walled, clear-bottom plate | **3917** | Corning |
| 78 | 384-well sterile, black-walled, clear-bottom plate | **3764** | Corning |

#### Table 2 - List of Antibodies

| S. No. | Antibody | Species | Catalog No. | Company | Dilution |
|---|---|---|---|---|---|
| 1 | CDK6 | Mouse | 3136 | Cell Signalling | 1:2000 (Western)<br>1:10 (Simple Western) |
| 2 | GFP | Rabbit | 2956 | Cell Signalling | 1:1000 (Western)<br>1:10 (Simple Western) |
| 3 | CRBN | Rabbit | 71810 | Cell Signalling | 1:1000 (Western)<br>1:100 (Simple Western) |
| 4 | VHL | Rabbit | 81292 | Cell Signalling | 1:10 |
| 5 | CDK4 | Rabbit | 12790 | Cell Signalling | 1:10 |
| 6 | HSP90 | Rabbit | 79641 | Cell Signalling | 1:1000 (Western)<br>1:50 (Simple Western) |
| 7 | Tubulin | Mouse | 3873 | Cell Signalling | 1:5000 (Western)<br>1:100 (Simple Western) |
| 8 | B-Actin | Rabbit | 4970 | Cell Signalling | 1:50 |
| 9 | GAPDH | Rabbit | 3683 | Cell Signalling | 1:50 |
| 10 | Vinculin | Rabbit | 13901 | Cell Signalling | 1:50 |
| 11 | pRb (807/811) | Rabbit | 8516 | Cell Signalling | 1:50 |
| 12 | pRb (780) | Rabbit | 9307 | Cell Signalling | 1:10 |
| 13 | Rb | Mouse | 9309 | Cell Signalling | 1:25 |
| 14 | Cyclin D1 | Rabbit | 2978 | Cell Signalling | 1:50 |
| 15 | Cyclin D3 | Mouse | 2936 | Cell Signalling | 1:10 |
| 16 | E2F1 | Rabbit | 3742 | Cell Signalling | 1:10 |
| 17 | CDK1 | Rabbit | 77055 | Cell Signalling | 1:10 |
| 18 | CDK2 | Rabbit | 2546 | Cell Signalling | 1:10 |
| 19 | Cyclin B1 | Rabbit | 4138 | Cell Signalling | 1:10 |
| 20 | Cyclin D2* | Rabbit | 3741 | Cell Signalling | 1:10 |
| 21 | Cyclin E1 | Rabbit | 20808 | Cell Signalling | 1:10 |
| 22 | p16* | Rabit | 18769 | Cell Signalling | 1:50 |
| 23 | p21 | Rabbit | 2947 | Cell Signalling | 1:50 |
| 24 | p27 | Rabbit | 3686 | Cell Signalling | 1:50 |
| 25 | p18* | Mouse | 2896 | Cell Signalling | 1:50 |
| 26 | Anti-Rabbit HRP | Goat | 3240 | Invitrogen | 1:2000 |
| 27 | Anti-Mouse | Goat | G21040 | Invitrogen | 1:5000 |
| 28 | Anti-Rabbit HRP | Goat | DM-001 | Simple Western | As is |
| 29 | Anti-Mouse HRP | Goat | DM-002 | Simple Western | As is |
| 30 | Streptavidin-HRP | | DM-004 | Simple Western | As is |

### Figures

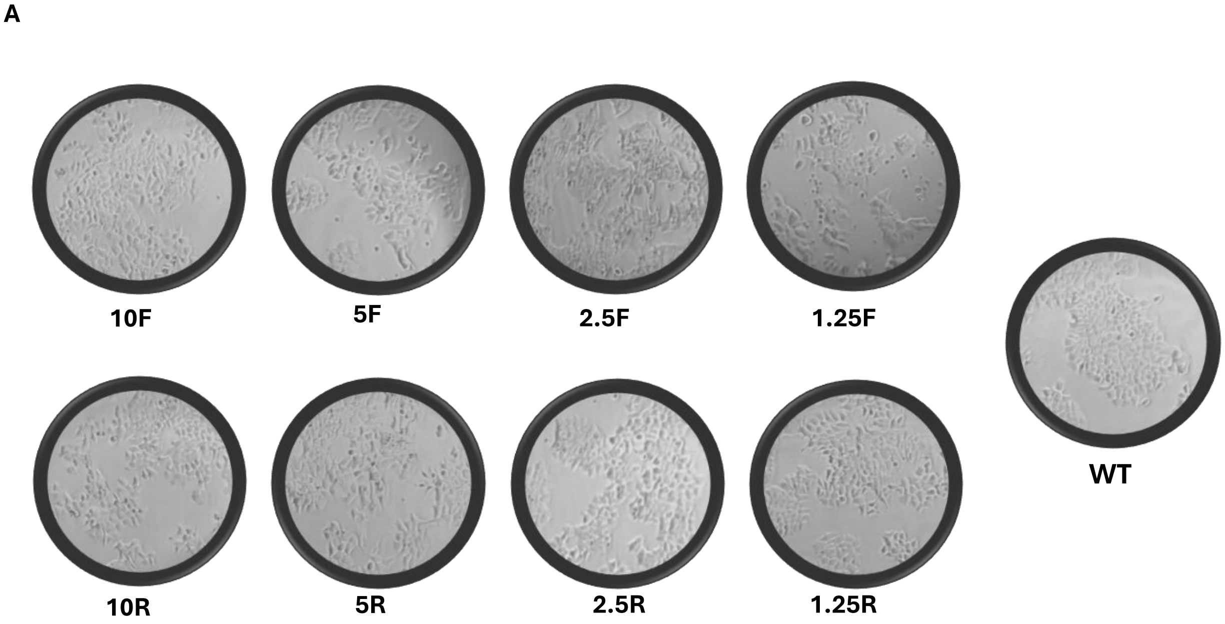


Figure S1 Validation MCF7-CDK6-GFP cell line A) Morphology comparison of the successfully transducted clones with MCF7 WT

B

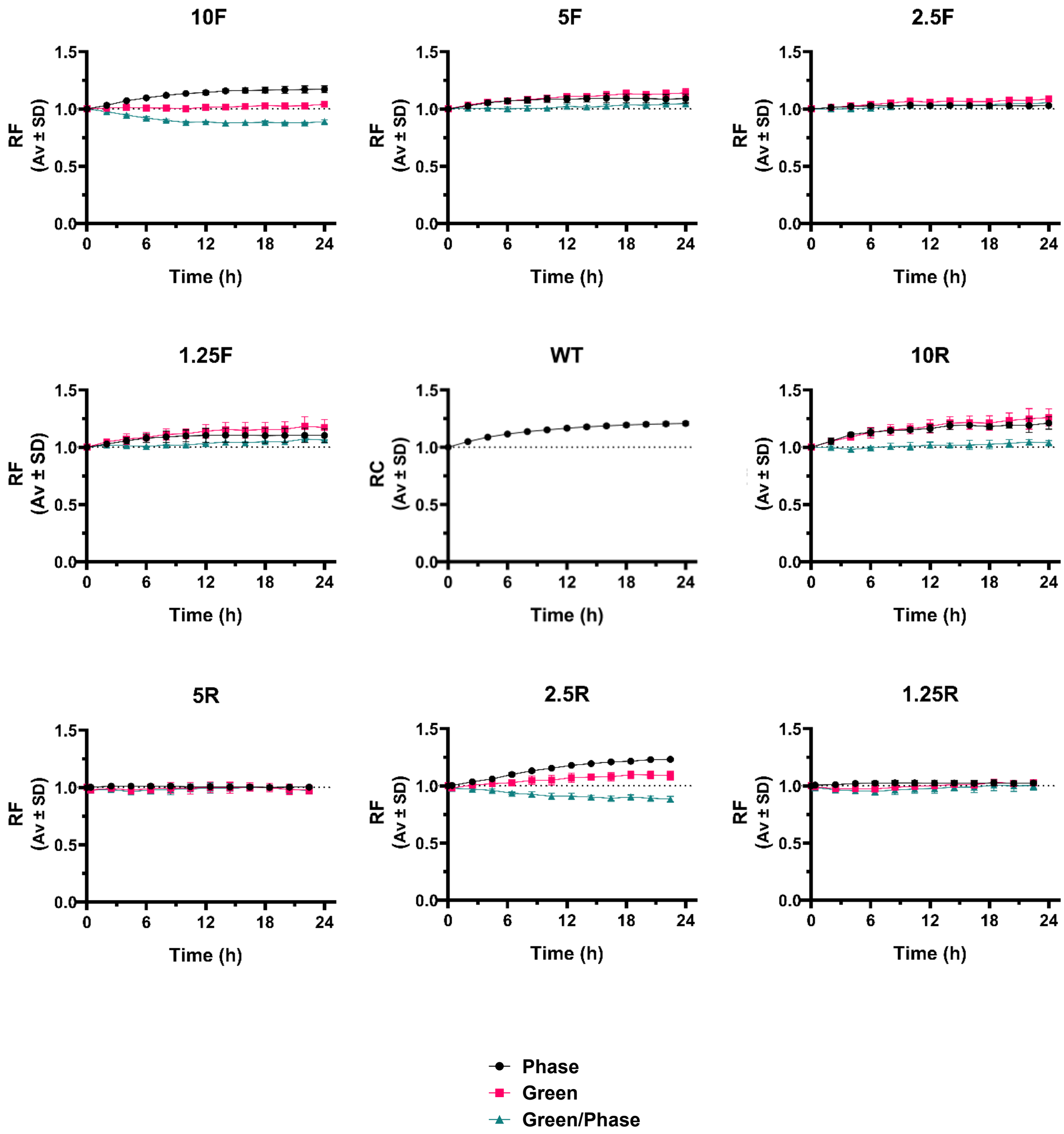


Figure S1 Validation MCF7-CDK6-GFP cell line contd.. B) 24-hour growth showing the phase confluence, green signal and the ratio of green/phase for all the MCF7-CDK6-GFP clones and MCF7 WT

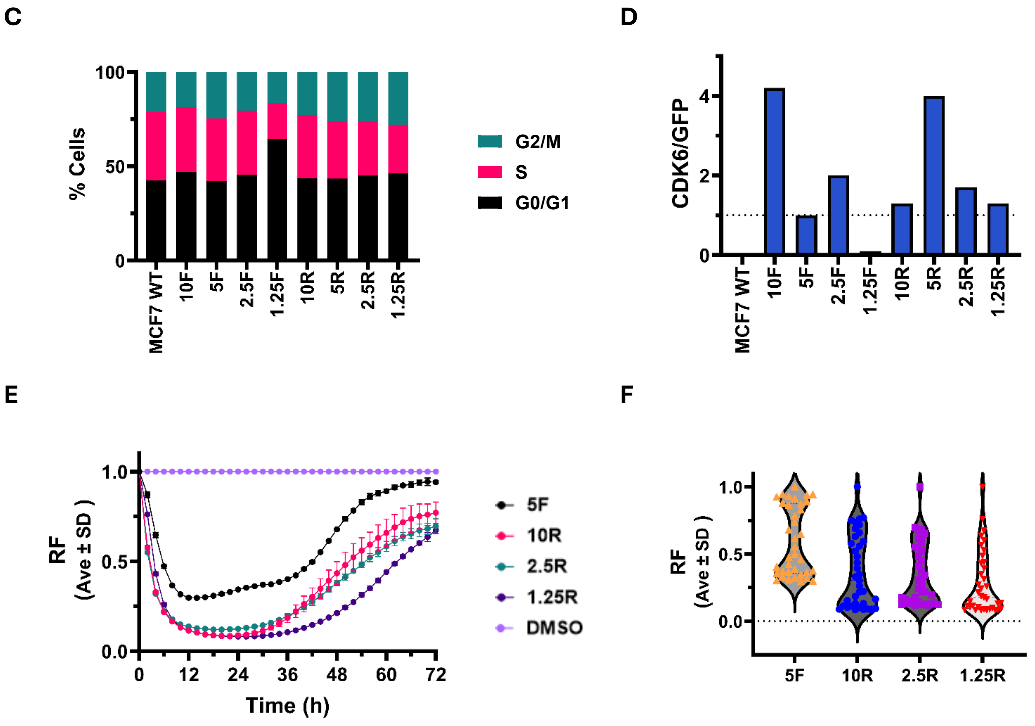


Figure S1 Validation MCF7-CDK6-GFP cell line contd.. C) Relative distribution of cell cycle phases for the MCF7-CDK6-GFP clones D) Ratio of CDK6 to GFP in the individual clones * *See Appendix A for blot images* E) Preliminary degradation experiment with 1 µM of BSJ-02-123 for 72 hours with the selected clones F) The average degradation of CDK6-GFP over 72 hours

G

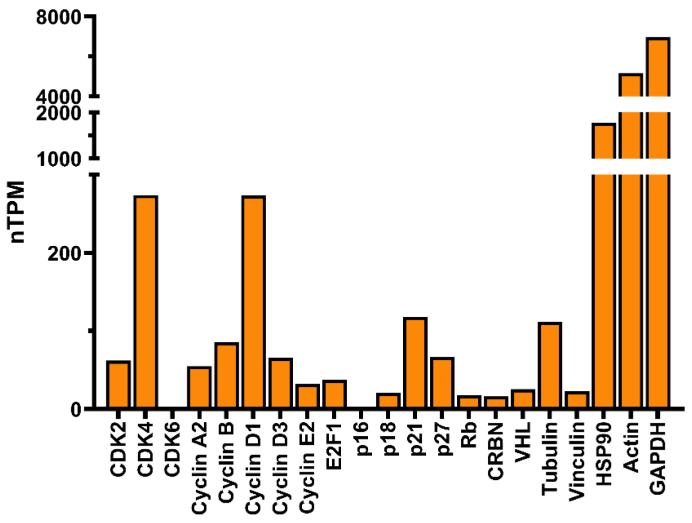


H

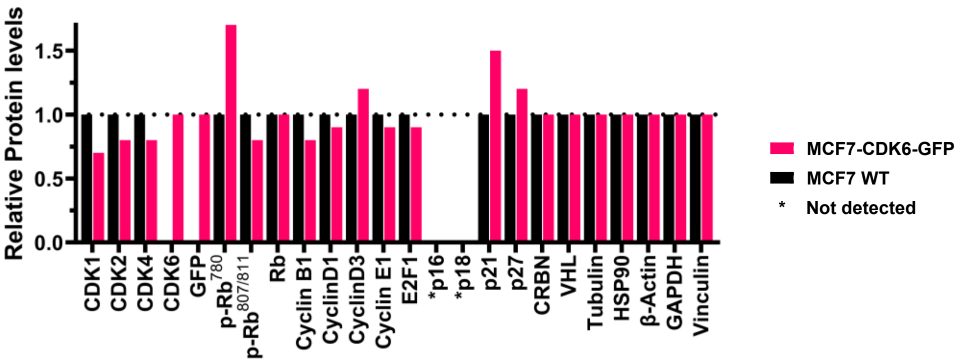


Figure S1 Validation MCF7-CDK6-GFP cell line contd.. G) Relative mRNA levels of cell cycle and housekeeping proteins per Protein Atlas H) Comparison of the cell cycle and housekeeping protein levels in MCF7WT and MCF7 CDK6-GFP clone 1.25R * *See Appendix B1 and B2 for blot images*

Figure S2 Time lapse video showing degradation of CDK6-GFP by DMSO or CDK6 PROTACs **videos digitally enhanced for contrast to aid visualization** ATTACHED separately

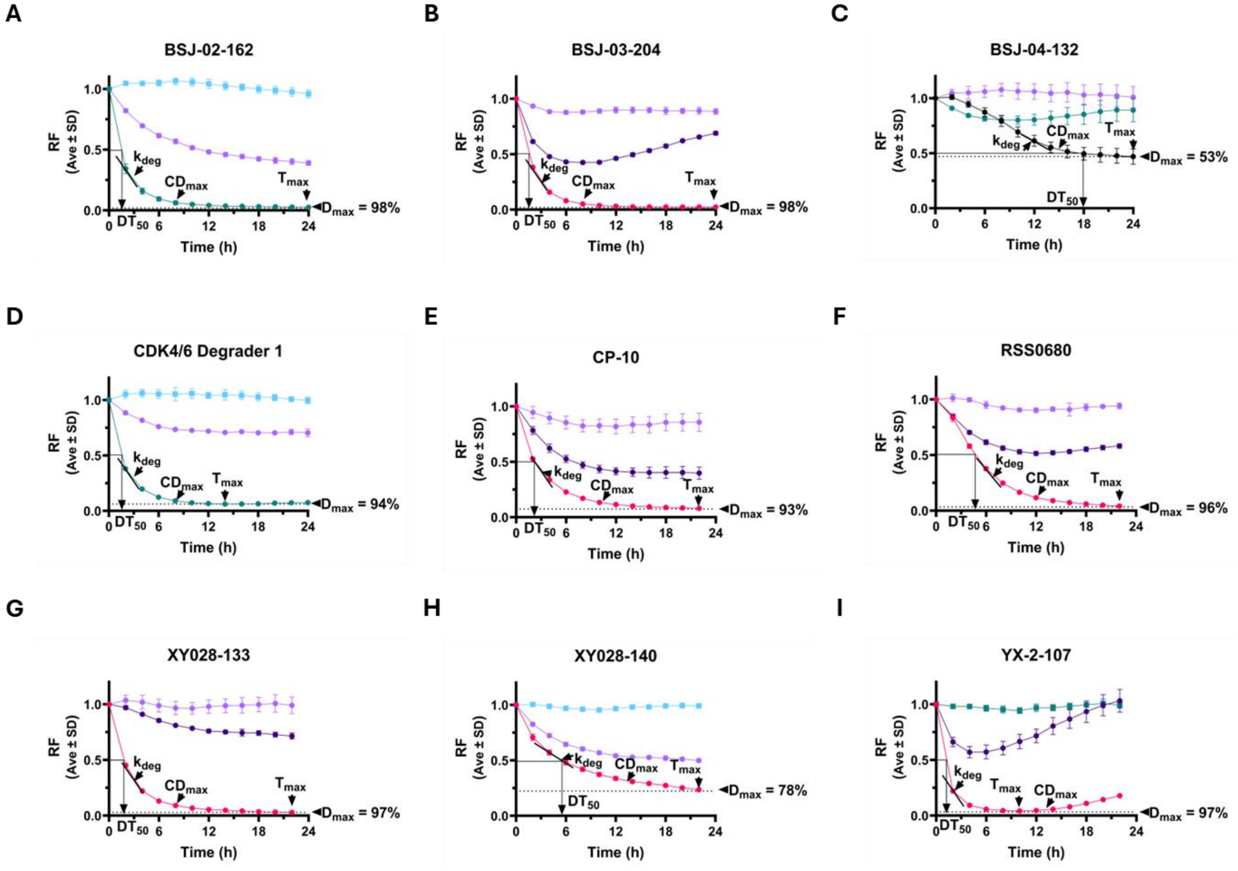


Figure S3 Degradation Plot showing parameters calculated - Maximum degradation ($D_{max}$) ;Concentration at which maximum degradation ($CD_{max}$ ); Time at which maximum degradation is observed ($T_{max}$) ; Time at which half of the PROTAC is degraded ($DT_{50}$); rate of degradation of the protein ($k_{deg}$) A) BSJ-02-162 B) BSJ-03-204 C) BSJ-04-132 D) CDK4/6 degrader 1 E) CP-10 F) RSS0680 G) XY028-133 H) XY028-140 I) YX-2-107

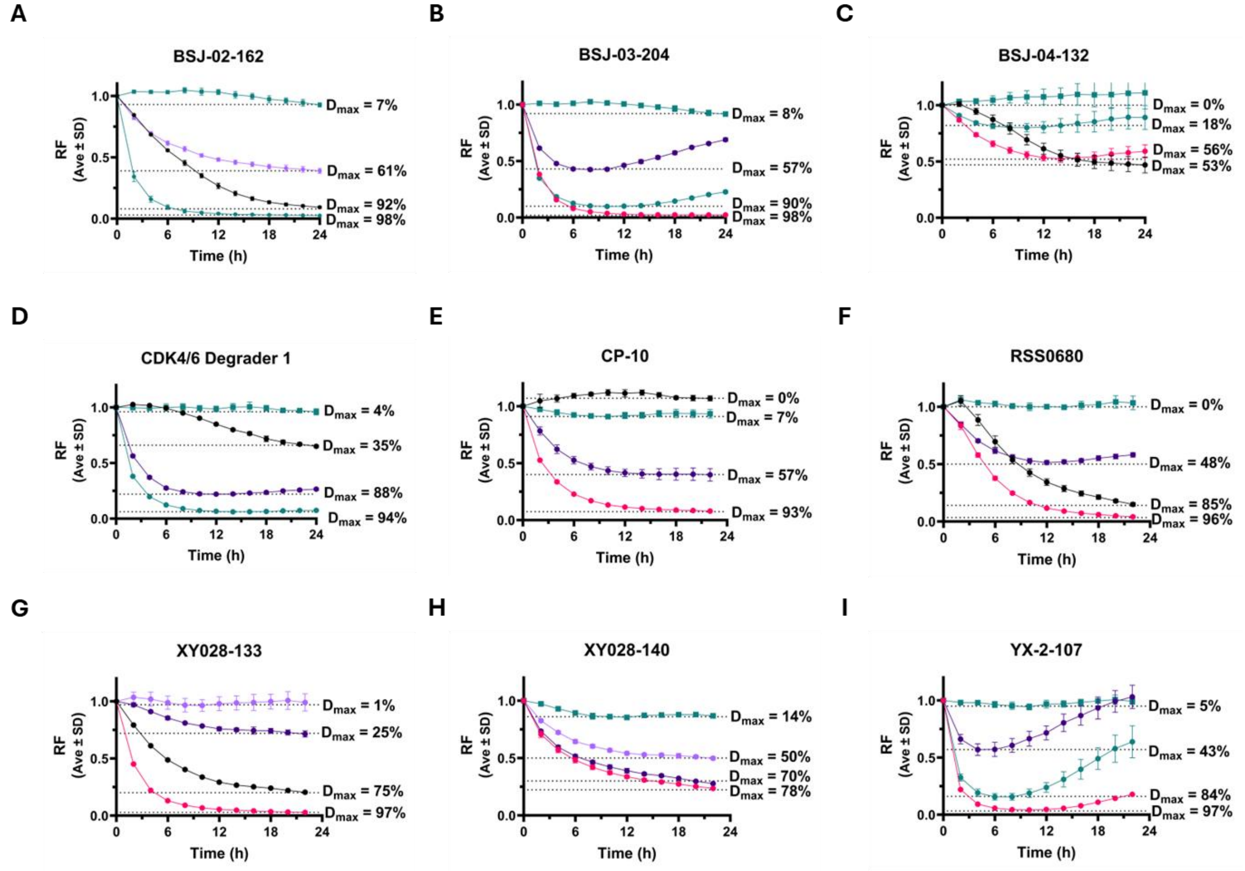


Figure S4 $D_{max}$ at various concentrations A) BSJ-02-162 B) BSJ-03-204 C) BSJ-04-132 D) CDK4/6 degrader 1 E) CP-10 F) RSS0680 G) XY028-133 H) XY028-140 I) YX-2-107

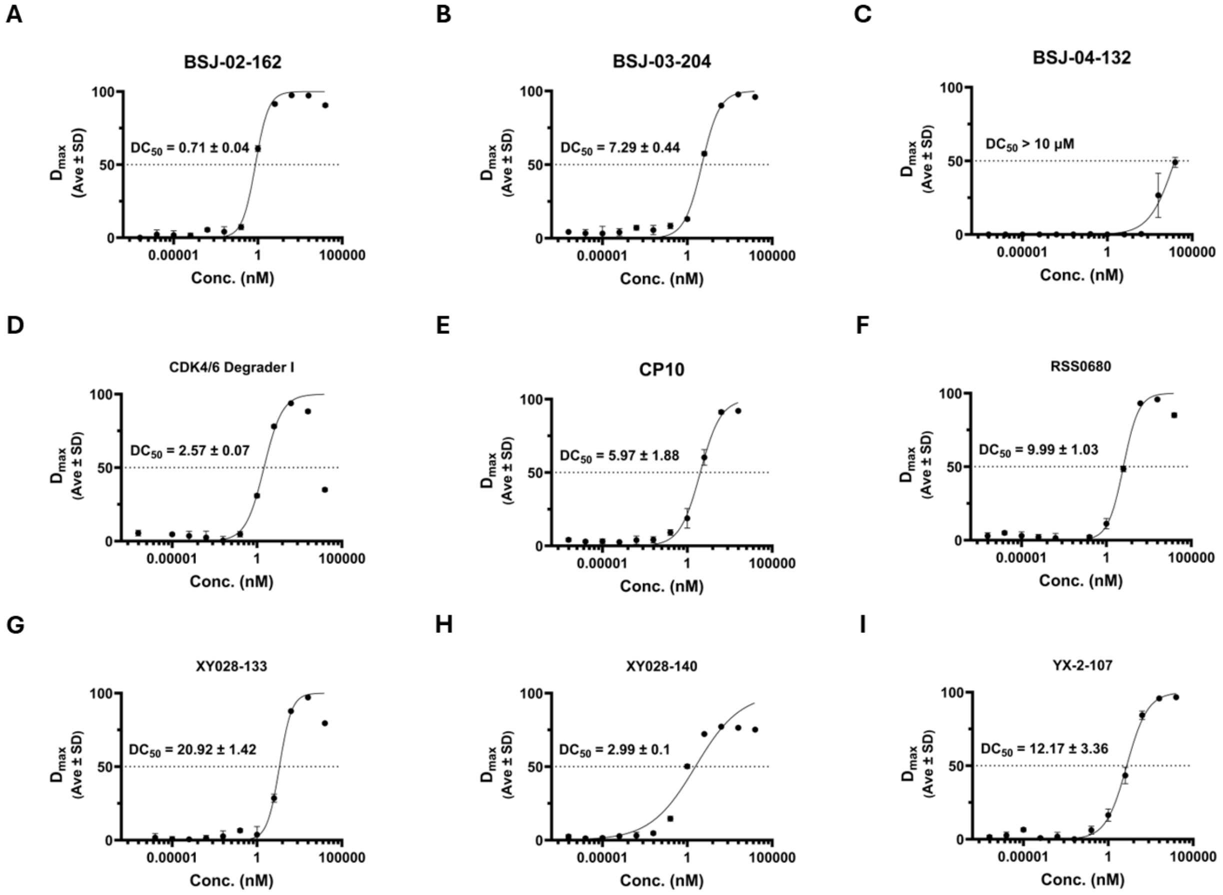


Figure S5 $DC_{50}$ for all PROTACS A) BSJ-02-162 B) BSJ-03-204 C) BSJ-04-132 D) CDK4/6 degrader 1 E) CP-10 F) RSS0680 G) XY028-133 H) XY028-140 I) YX-2-107

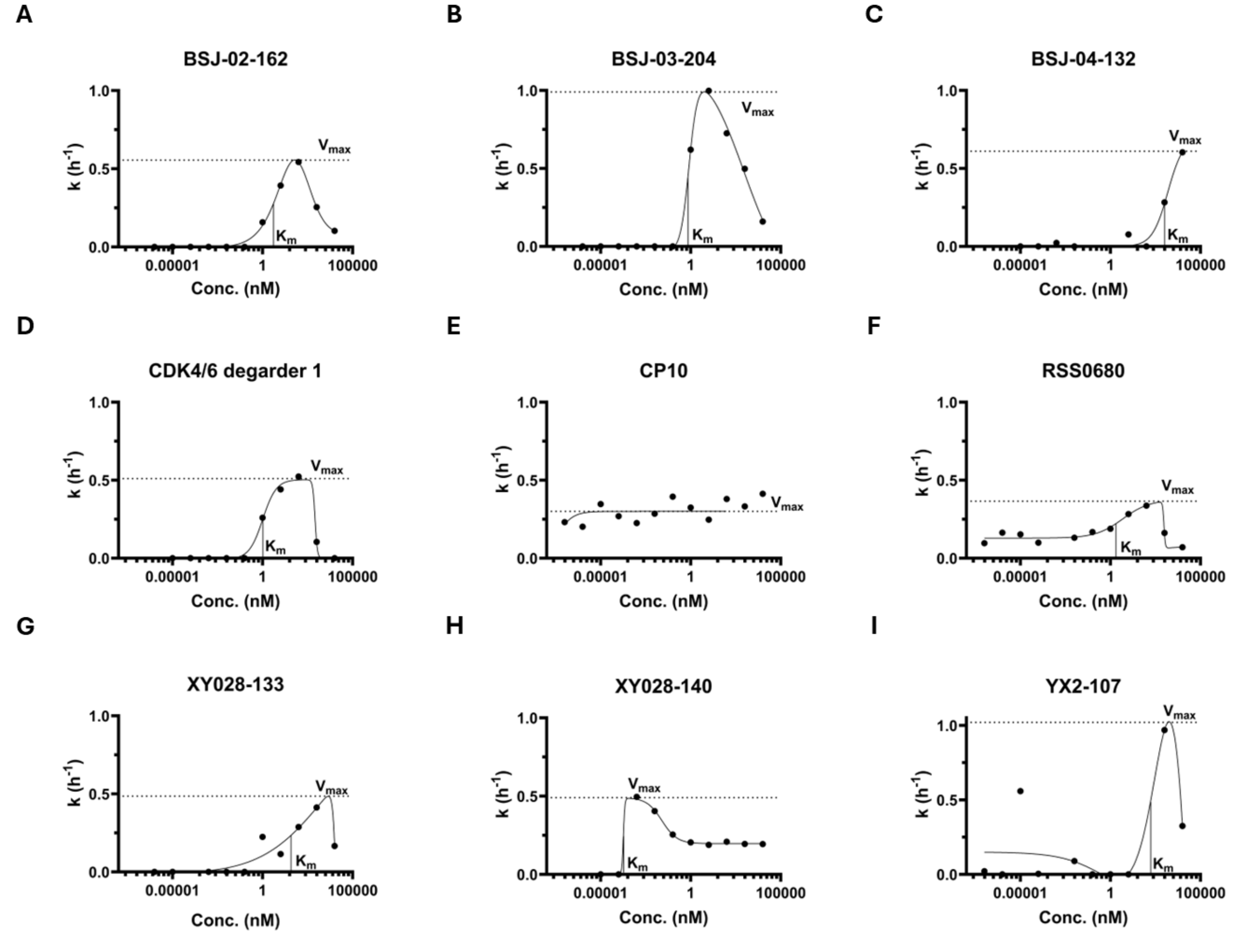


Figure S6 $V_{max}$ and $K_m$ of each PROTAC A) BSJ-02-162 B) BSJ-03-204 C) BSJ-04-132 D) CDK4/6 degrader 1 E) CP-10 F) RSS0680 G) XY028-133 H) XY028-140 I) YX-2-107

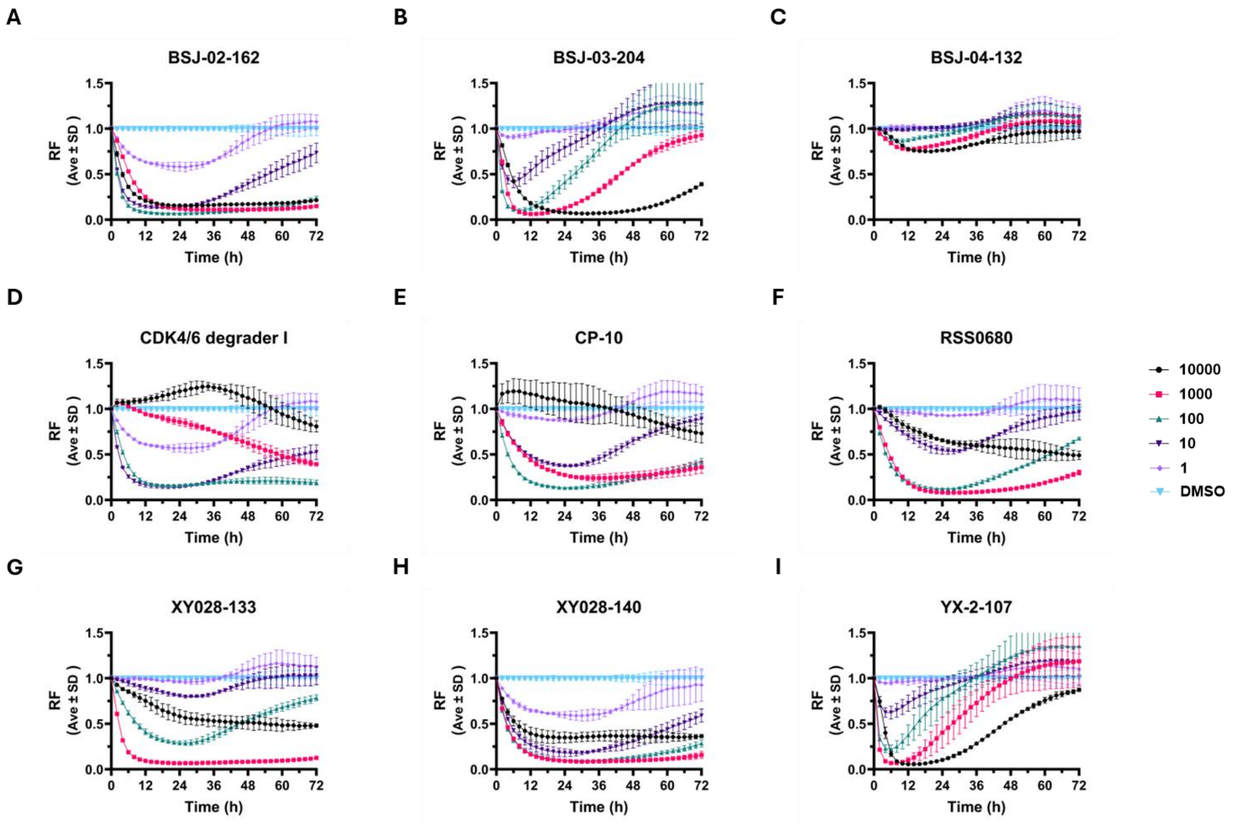


Figure S7 72-hour dose response curves for all PROTAC A) BSJ-02-162 B) BSJ-03-204 C) BSJ-04-132 D) CDK4/6 degrader 1 E) CP-10 F) RSS0680 G) XY028-133 H) XY028-140 I) YX-2-107

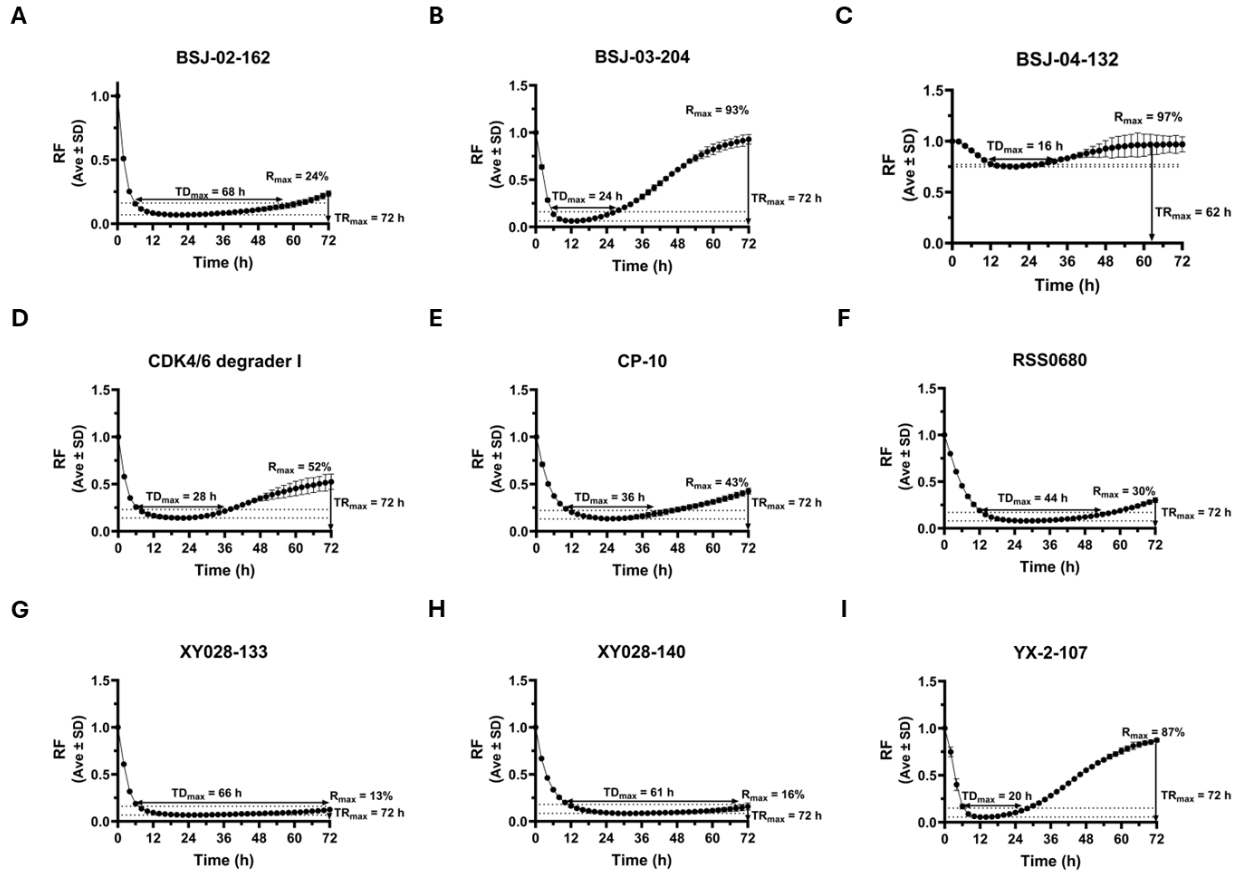


Figure S8 Plots showing $TD_{max}$, $R_{max}$, and $TR_{max}$ for all PROTACs A) BSJ-02-162 B) BSJ-03-204 C) BSJ-04-132 D) CDK4/6 degrader 1 E) CP-10 F) RSS0680 G) XY028-133 H) XY028-140 I) YX-2-107

A

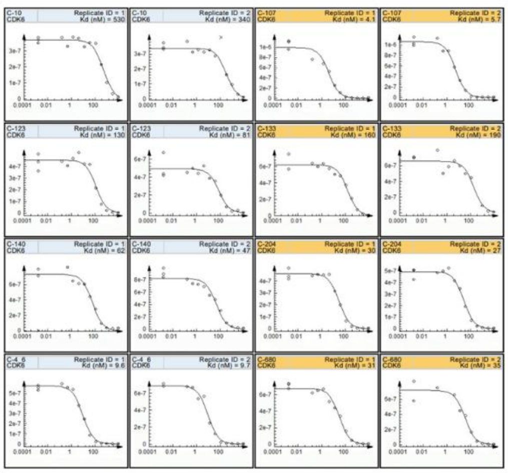

B

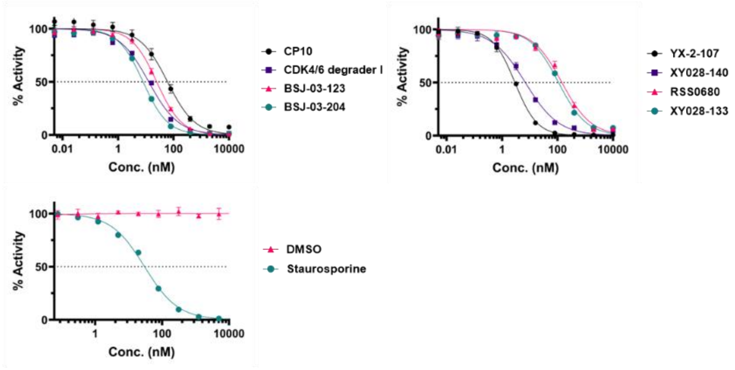


C

| PROTAC | $IC_{50}$ (nM) | $K_d$ (nM) |
|---|---|---|
| BSJ-03-123 | 25.72 ± 16.3 | 105.5 ± 34.65 |
| BSJ-03-204 | 8.58 ± 0.43 | 285.5 ± 2.12 |
| CDK4/6 Degrader I | 15.31 ± 1 | 9.65 ± 0.07 |
| CP10 | 50.59 ± 6.33 | 435.0 ± 134.35 |
| RSS0680 | 167.63 ± 22.27 | 33.0 ± 2.83 |
| XY028-133 | 124.43 ± 7.5 | 175.0 ± 21.21 |
| XY028-140 | 7.73 ± 0.63 | 54.5 ± 10.61 |
| YX-2-107 | 2.98 ± 0.02 | 4.9 ± 1.13 |

Figure S9 Biophysical assays A) Binding assay - Kd values from a KdELECT screen performed by Eurofins C-10 (CP-10); C-107 (YX-2-107); C-123 (BSJ-03-123); C-133 (XY028-133); C-140 (XY028-140); C-204 (BSJ-03-204); C-4-6 (CDK4/6 Degrader I) and C-680 (RSS0680) B) Activity assay - $IC_{50}$ curves from a HotSpot Kinase assay performed by Reaction Biology C) Table of $K_d^{binary}$ and $IC_{50}$ calculated from these assays

A

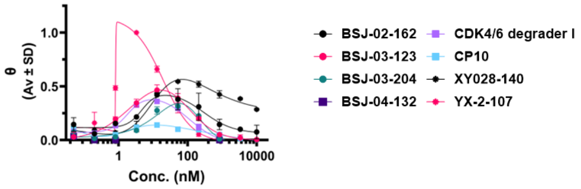


H

| PROTAC | $TF_{50}$ | $TI_{50}$ | CW | AUC | $TC_{max}$ |
|---|---|---|---|---|---|
| | *(nM)* | *(nM)* | *(nM)* | | *(nM)* |
| BSJ-02-162 | 3.73 ± 1.57 | 375.35 ± 133.7 | 371.709 | 1105 | 20 |
| BSJ-03-123 | 0.96 ± 0.06 | 145.85 ± 9.12 | 144.88945 | 316 | 15 |
| BSJ-03-204 | 7.51 ± 0.44 | 294.2 ± 10.96 | 286.6875 | 156 | 70 |
| BSJ-04-132 | ND | ND | ND | ND | ND |
| CDK4/6 Degrader I | 1.31 ± 0.13 | 157.05 ± 25.53 | 155.7395 | 273 | 10 |
| CP10 | 1.74 | 521.35 ± 28.07 | 519.61 | 190 | 10 |
| XY028-140 | 10.17 ± 1.41 | 13888.50 ± 826.61 | 13878.329 | 3647 | 100 |
| YX-2-107 | 0.79 ± 0.01 | 26.15 ± 2.21 | 25.35235 | 198 | 4 |

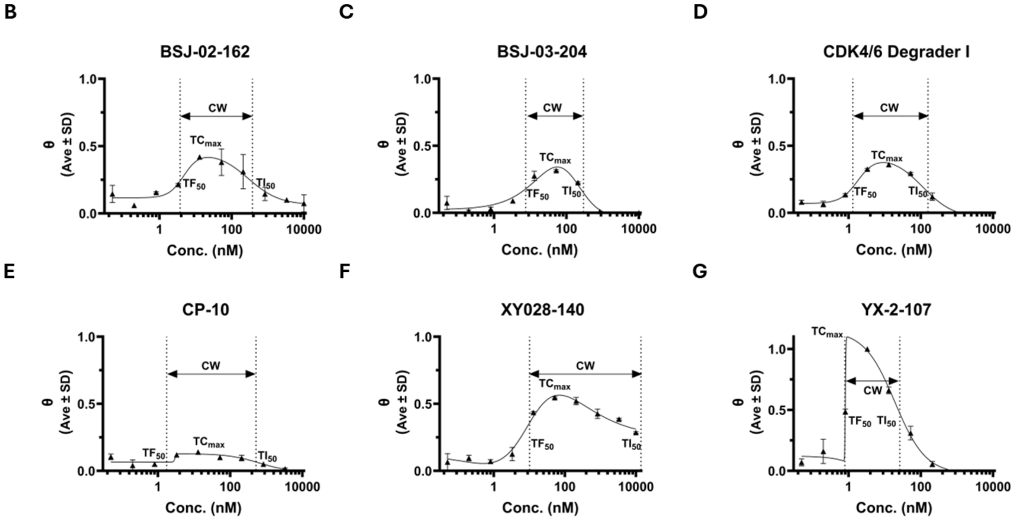


Figure S10 AlphaLISA A) Fraction bound curve for all PROTACs and representative curves showing half maximum formation of TC ($TF_{50}$); half maximum inhibition of TC ($TI_{50}$); Concentration at which maximum TC formation is seen is $TC_{max}$ and the concentration window (CW) where TC formation is favorable B) BSJ-02-162 C) BSJ-03-204 D) CDK4/6 degrader 1 E) CP-10 F) XY028-140 G) YX-2-107 H) Parameters calculated from an Alpha curve

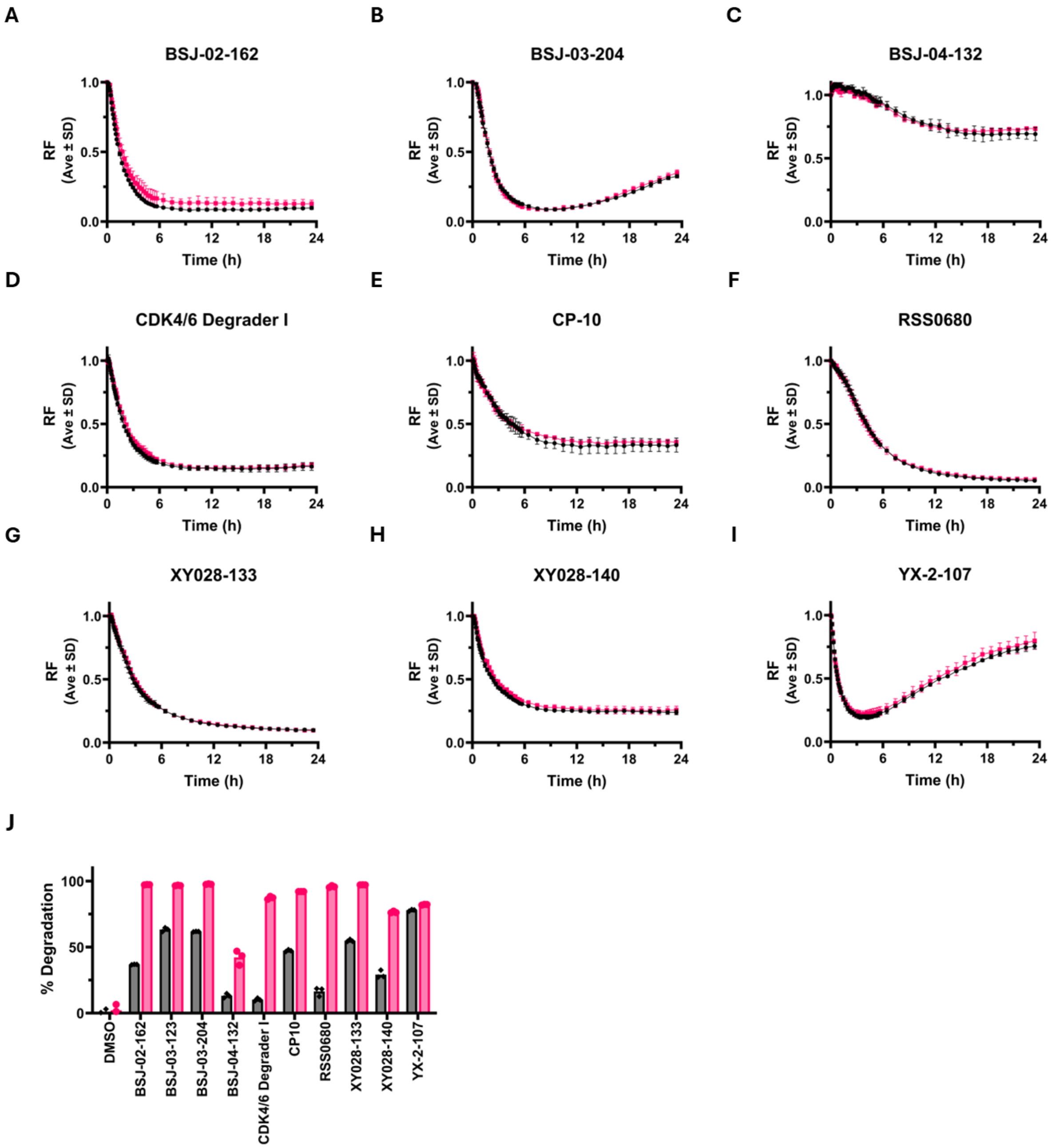


Figure S11 Permeability of all PROTAC's at CDmax concentration A) BSJ-02-162 B) BSJ-03-204 C) BSJ-04-132 D) CDK4/6 degrader 1 E) CP-10 F) RSS0680 G) XY028-133 H) XY028-140 I) YX-2-107 J) Comparison of the 2-hr and 22-hour degradation of CDK6-GFP

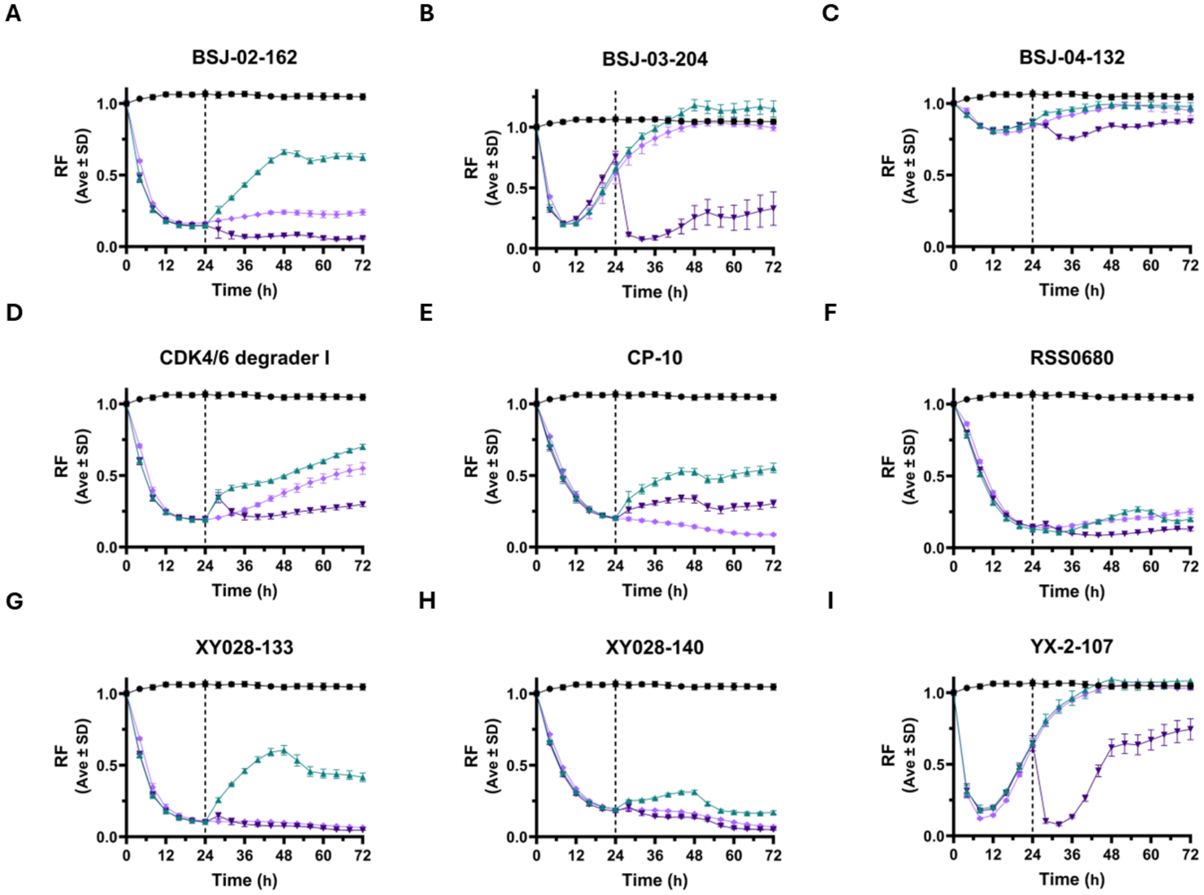


Figure S12 Washout and retreat for all PROTAC's at CDmax concentration A) BSJ-02-162 B) BSJ-03-204 C) BSJ-04-132 D) CDK4/6 degrader 1 E) CP-10 F) RSS0680 G) XY028-133 H) XY028-140 I) YX-2-107

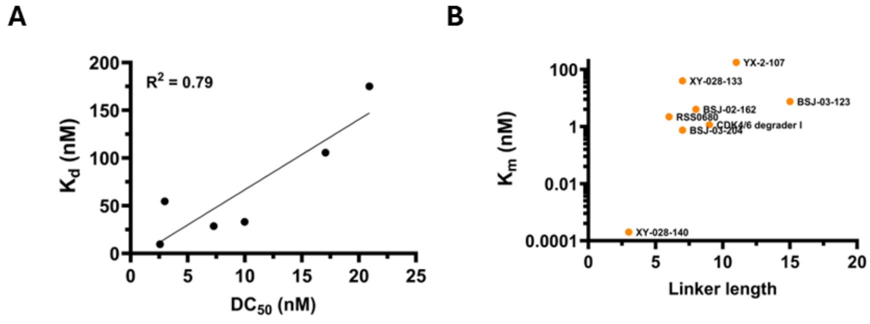


Figure S13 Potency Correlation plots A) Correlation of affinity and potency B) Comparison of linker length and Km

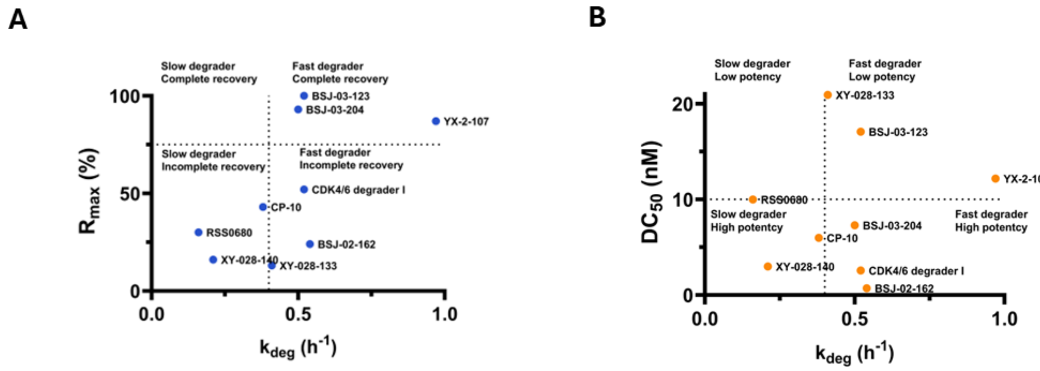


Figure S14 Classification plots A) Comparison of efficacy and efficiency B) Comparison of potency and efficiency

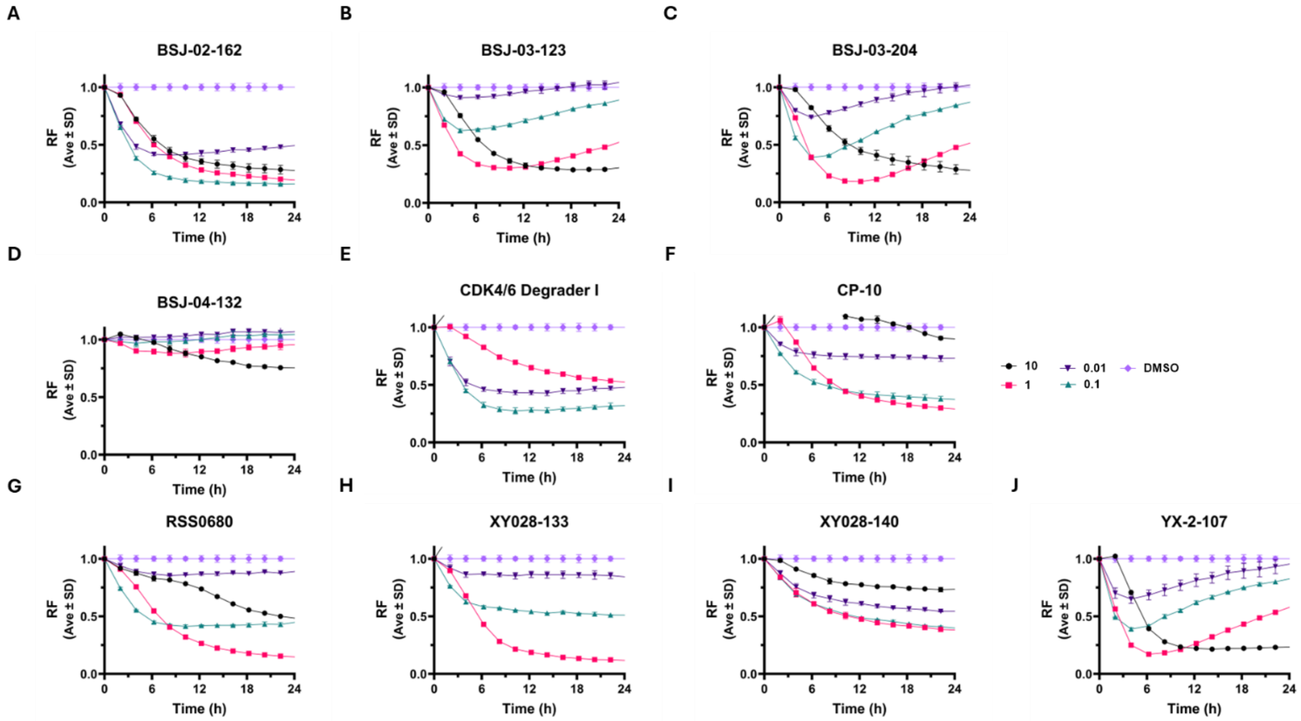


Figure S15 Miniaturization – Dose response curves done in a 384-well plate A) BSJ-02-162 B) BSJ-03-123 C) BSJ-03-204 D) BSJ-04-132 E) CDK4/6 degrader 1 F) CP-10 G) RSS0680 H) XY028-133 I) XY028-140 J) YX-2-107

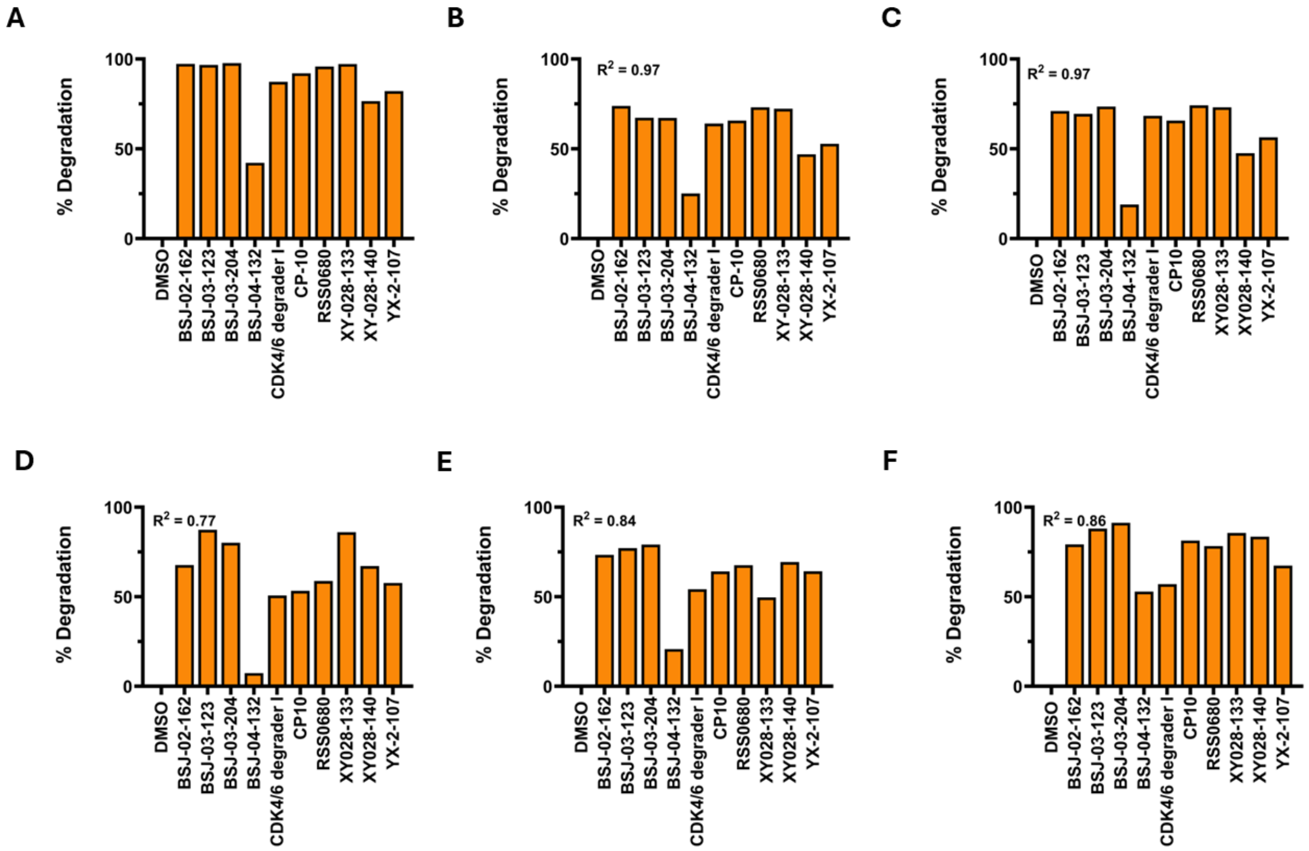


**G**

| PROTAC | Rank | | | | |
|---|---|---|---|---|---|
| | *FL* | *TR-FRET* | *WB* | *SW* | *SW BxPC3* |
| BSJ-02-162 | 1 | 1 | 4 | 3 | 6 |
| BSJ-03-123 | 5 | 4 | 1 | 2 | 2 |
| BSJ-03-204 | 4 | 3 | 3 | 1 | 1 |
| BSJ-04-132 | 10 | 10 | 10 | 10 | 10 |
| CDK4/6 Degrader I | 6 | 6 | 9 | 8 | 9 |
| CP10 | 7 | 7 | 8 | 7 | 5 |
| RSS0680 | 2 | 2 | 6 | 5 | 7 |
| XY028-133 | 3 | 5 | 2 | 9 | 3 |
| XY028-140 | 9 | 9 | 5 | 4 | 4 |
| YX-2-107 | 8 | 8 | 7 | 6 | 8 |

**H**

| PROTAC | $GI_{50}$ (*μM*) |
|---|---|
| BSJ-02-162 | 2.49 +/- 1.69 |
| BSJ-03-123 | >10 |
| BSJ-03-204 | >10 |
| BSJ-04-132 | >10 |
| CDK4/6 Degrader I | 0.019 +/- 0.006 |
| CP10 | >10 |
| RSS0680 | 0.35 +/- 0.028 |
| XY028-133 | 1.93 +/- 0.28 |
| XY028-140 | >10 |
| YX-2-107 | 3.88 +/- 0.36 |

Figure S16 Comparison of degradation seen on Incucyte A) with orthogonal assays performed with lysates obtained from 1 µM 24-hour PROTAC treatment for B) Fluorescence C) TR-FRET Lanthascreen D) Traditional Western blot * *See Appendix C for blot images* E) Simple Western with MCF7-CDK6-GFP* *See Appendix D for blot images* F) Simple Western with BxPC3 * *See Appendix E for blot images* G) Rankings based on orthogonal assays H) $GI_{50}$ from a 120-hour assay
